# Valence and salience encoding by parallel circuits from the paraventricular thalamus to the nucleus accumbens

**DOI:** 10.1101/2023.07.03.547570

**Authors:** Jean K. Rivera-Irizarry, Peter U. Hámor, Sydney A. Rowson, Joseph Asfouri, Dezhi Liu, Lia J. Zallar, Aaron F. Garcia, Mary Jane Skelly, Kristen E. Pleil

## Abstract

The anterior and posterior subregions of the paraventricular thalamus (aPVT and pPVT, respectively) play unique roles in learned behaviors, from fear conditioning to alcohol/drug intake, potentially through differentially organized projections to limbic brain regions including the nucleus accumbens medial shell (mNAcSh). Here, we found that the aPVT projects broadly to the mNAcSh and that the aPVT-mNAcSh circuit encodes positive valence, such that *in vivo* manipulations of the circuit modulated both innately programmed and learned behavioral responses to positively and negatively valenced stimuli, particularly in females. Further, the endogenous activity of aPVT presynaptic terminals in the mNAcSh was greater in response to positively than negatively valenced stimuli, and the probability of synaptic glutamate release from aPVT neurons in the mNAcSh was higher in females than males. In contrast, we found that the pPVT-mNAcSh circuit encodes stimulus salience regardless of valence. While pPVT-mNAcSh circuit inhibition suppressed behavioral responses in both sexes, circuit activation increased behavioral responses to stimuli only in males. Our results point to circuit-specific stimulus feature encoding by parallel PVT-mNAcSh circuits that have sex-dependent biases in organization and function.

---

Performing the most appropriate behavior in response to an environmental stimulus is a core feature of adaptive behavior, and this action requires accurate assessment and integration of stimulus salience and valence^1–7^. Plasticity in circuits encoding these stimulus features may contribute to maladaptive behaviors such as aberrant hedonic value attribution or anhedonia^2, 7–9^, phenotypes commonly expressed across neuropsychiatric diseases including alcohol/substance use and affective disorders^7–12^. Males and females are differentially susceptible to these diseases^13–18^, and they exhibit both basic and disease-associated differences in reward and aversion behaviors that may be due to sex-dependent circuit organization and function^13, 16, 19–24^. The paraventricular nucleus of the thalamus (PVT) is a midline thalamic brain region highly interconnected with the limbic system via glutamatergic projection neurons that regulates a myriad of stress and reward-related behaviors from natural and drug reward seeking to conditioned fear^25–36^, and recent evidence suggests that sex-dependent function of excitatory PVT circuits plays a role in behavioral sex differences^24^.

The PVT participates in guiding the motivated behavioral responses to positively and negatively valenced conditioned stimuli^25–27, 29, 37^, including when these are in direct conflict^5, 6^. Studies (almost exclusively in males) show that the glutamatergic projection from the PVT to the nucleus accumbens (NAc), particularly the medial shell subregion (mNAcSh), may be a primary circuit hub for these roles; however, they provide conflicting evidence regarding the role of the PVT-NAc circuit in regulating behavior, with some showing the activity of this circuit promotes natural and drug reward-seeking behaviors, while others show it is aversive, involved in fear, or negatively associated with appetitive reward behavior^5, 6, 25, 26, 29, 35, 38–40^. These seemingly contradictory findings for the PVT-NAc circuit’s roles in learned behaviors may be related to the unique functions and circuit organization of the anterior vs. posterior PVT (aPVT and pPVT, respectively)^5, 6, 34, 40^. The aPVT has generally been shown to be involved in natural and drug reward-related behaviors while the pPVT is involved in aversive behaviors, fear conditioning, and chronic stress^29, 41^. However, neurons in both the aPVT and pPVT respond to positively and negatively valenced stimuli and their cues during conditioning^5, 6, 32, 34^ and display unique combinatorial responses to these stimuli depending on their valence and salience^5, 6, 32, 34^.

Given that glutamatergic synaptic transmission in the mNAcSh modulates both reward and aversion behaviors^42, 43^, we posited that the aPVT and pPVT provide specifically organized synaptic inputs to the mNAcSh to encode valence and salience information required for adaptive behavioral responses to both appetitive and aversive environmental stimuli. Using anatomical tracing, slice electrophysiology, and *in vivo* chemogenetic manipulations and biosensor imaging during behavior, here we show that parallel PVT-mNAcSh circuits are essential for behavioral responses to salient stimuli with positive and negative valence. We found that the aPVT-mNAcSh and pPVT projections to the mNAcSh participate in valence and salience encoding, respectively, playing essential, complementary roles in innately programmed behavioral responses to environmental stimuli; these functions further extend to behavioral responses to conditioned stimuli following associative learning. We further found that the organization and function of these PVT-mNAcSh circuits are more complex than previously appreciated, including notable sexual dimorphisms in the strength of synaptic connections and glutamatergic efficiency at aPVT-mNAcSh synapses and the necessity and sufficiency of aPVT- and pPVT-mNAcSh circuit activity to regulate stimulus encoding. These results have important implications for sex differences in adaptive behaviors and the circuits underlying them, and they resolve current ambiguity in the literature about the role of PVT-NAc circuitry in behavior.

## Results

### Sexual dimorphism in glutamatergic efficiency at aPVT-mNAcSh synapses

We performed retrograde tracing to characterize the extent to which neurons across the anterior-posterior (A-P) extent of the PVT project to the mNAcSh (**Fig. 1a,b**). In both sexes, the proportion of DAPI+ PVT cells that project to the mNAcSh was robust across the entire PVT, with a bias toward the aPVT compared to pPVT (**Fig. 1c,d**). Analysis of the distribution of the retrogradely labeled population of mNAcSh-projecting PVT neurons across the A-P axis of the PVT within individual mice ((# retrogradely labeled neurons at A-P coordinate) / (total # retrogradely labeled neurons across PVT)) confirmed a bias toward the aPVT (**Fig. 1e**). These anatomical tracing results suggest that the aPVT displays a particularly dense projection to the mNAcSh in both males and females. Therefore, we used an *ex vivo* optogenetic + slice electrophysiology approach to measure the function of the aPVT-mNAcSh pathway and evaluate whether there were differences in the functional synaptic innervation and strength of the dorsal vs. ventral mNAcSh by aPVT inputs (**Fig. 2**), given some evidence that these mNAcSh subregions may play opposing behavioral roles^44^. In mice injected with channelrhodopsin (ChR2) in the aPVT (**Fig. 2a; Fig. S1**), we found that optically-evoked excitatory postsynaptic currents (oEPSCs) in mNAcSh neurons, a measure of glutamatergic synaptic transmission between aPVT presynaptic terminals and mNAcSh postsynaptic neurons, were similar in magnitude in dorsal and ventral mNAcSh neurons and between sexes (**Fig. 2b,c**). To control for possible differences in viral expression between mice and directly compare the input density to the dorsal vs. ventral mNAcSh, we assessed oEPSC magnitude in paired neurons within the same slice and calculated the relative ratio of the dorsal oEPSC amplitude compared to ventral oEPSC: oEPSC_Dorsal_ / (oEPSC_Dorsal_ + oEPSC_Ventral_). This ratio was approximately 0.50 in both male and female mice (**Fig. 2d**), confirming that the aPVT similarly innervates the dorsal and ventral mNAcSh within individual mice (and specifically within mNAcSh Bregma coordinates), as well as across sex. While the magnitude of oEPSCs was similar between males and females, the paired pulse ratio (PPR) measured from aPVT terminal oEPSCs in both mNAcSh subregions was higher in males than females and above 1.0 in males but not females (**Fig. 2e**). These data suggest that glutamatergic efficiency of aPVT-mNAcSh synapses related to short-term plasticity is higher in females than males. Specifically, these results suggest that the probability of synaptic glutamate release from aPVT terminals in the mNAcSh is low in males but high in females, suggesting this neuron population acts as a high-pass filter in males, requiring relatively more robust stimulation to elicit glutamate release than females.

**Fig. 1:**
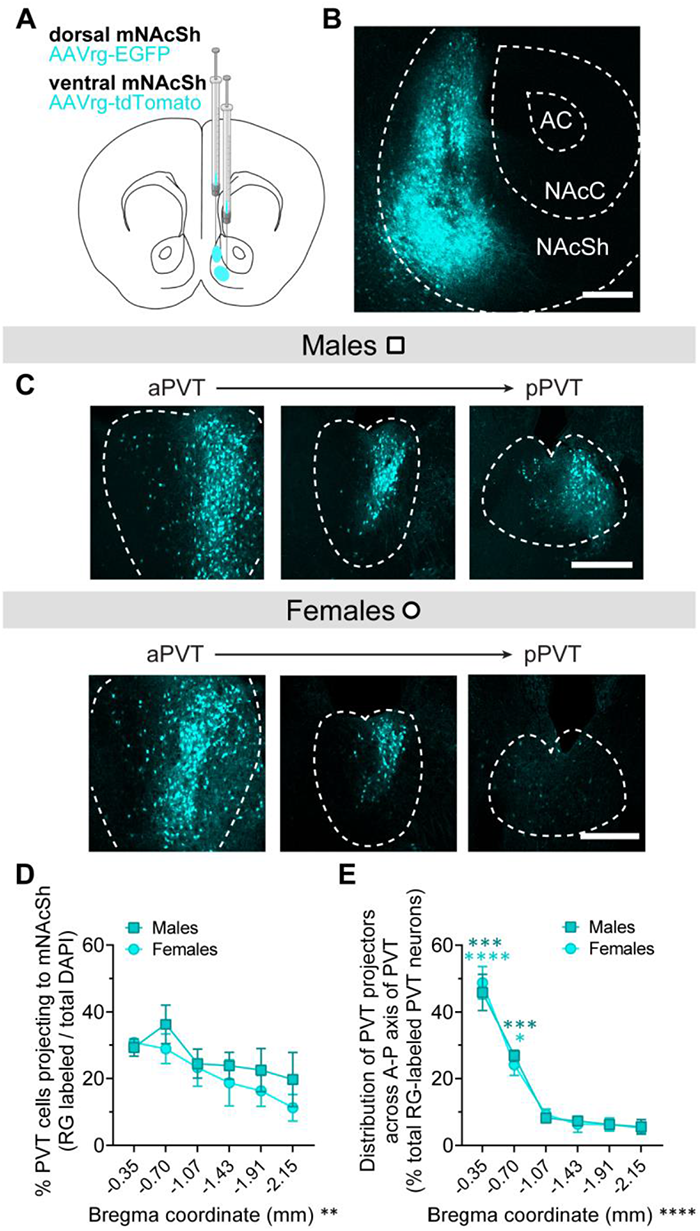
Sex differences in the topographical organization of PVT-mNAcSh projections. **a)** Schematic of the viral strategy to retrogradely label neurons projecting to the mNAcSh. **b)** Representative image of NAc showing retrograde tracer expression at the site of injection in unilateral mNAcSh. Scale bar = 250 µm. **c)** Representative images across the anterior-posterior extent of the PVT in a male (top) and female (bottom) showing retrogradely labeled mNAcSh projection neurons. Scale bar = 500 µm. **d)** % of PVT cells that project to the mNAcSh across the A-P extent of the PVT: (# retrogradely labeled neurons) / (total DAPI+). **e)** Distribution of mNAcSh-projecting neurons across the A-P extent of the PVT within individual mice: (# retrogradely labeled neurons at A-P coordinate) / (total # retrogradely labeled neurons across PVT). \*\**P* < 0.01, \*\*\*\**P* < 0.0001 for main effects of Bregma coordinate.

**Fig. 2:**
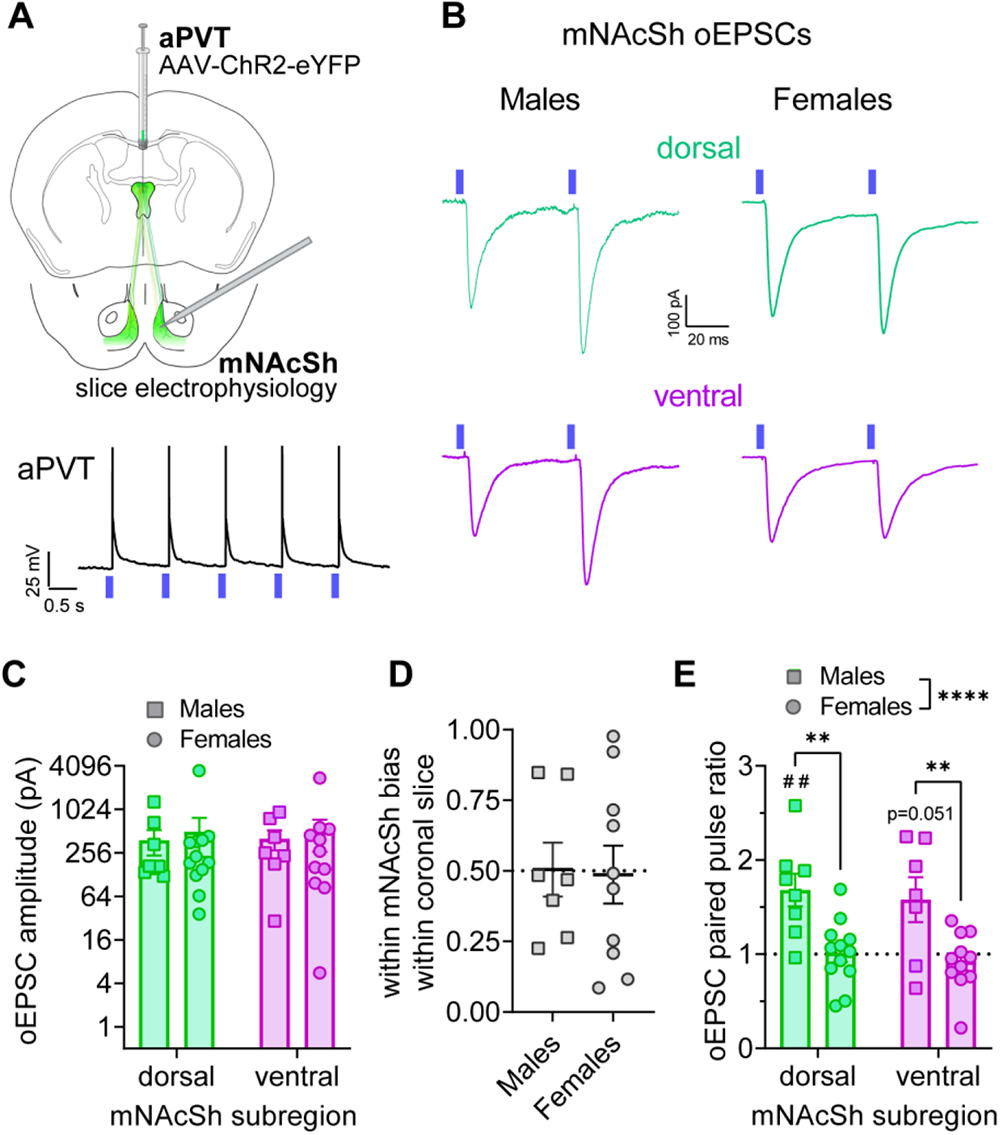
aPVT neurons exhibit greater probability of synaptic glutamate release in the mNAcSh in females than males. **a)** Top: Schematic of strategy to express ChR2 in the aPVT-mNAcSh circuit and perform *ex vivo* slice electrophysiology recordings in the mNAcSh. Bottom: Representative traces of 470 nm LED optically-elicited action potentials aPVT cell bodies during current-clamp recordings used to functionally confirm ChR2 expression in aPVT neurons. **b)** Representative traces of optically-evoked excitatory postsynaptic currents (oEPSCs) in neurons in the dorsal (green) and ventral (purple) mNAcSh in response to ChR2-elicited glutamate release from aPVT presynaptic terminals in males (left) and females (right). **c)** Amplitude of oEPSCs in dorsal and ventral mNAcSh neurons. **d)** Bias in oEPSC amplitude toward the dorsal mNAcSh, measured by the relative ratio of oEPSC amplitude in the dorsal mNAcSh compared to ventral mNAcSh within individual *ex vivo* slices (amplitude^dorsal^ / (amplitude^dorsal^ + amplitude^ventral^). **e)** The paired pulse ratio (PPR) of oEPSCs (amplitude^pulse^ ^2^ / amplitude^pulse^ ^1^) in dorsal and ventral mNAcSh neurons is higher in males than females and above 1.0 in males. \*\**P* < 0.01, \*\*\*\**P* < 0.0001 for ANOVA main effect and post hoc unpaired t-tests between groups within subregion as indicated; ^##^*P* < 0.01 for one-sample t-tests compared to 1.0 within each group.

### aPVT-mNAcSh circuit activity promotes stimulus valence

Given our anatomical and ChR2-assisted functional mapping results showed that there are robust projections to the mNAcSh across the PVT that are anatomically similar between sexes but functionally more robust in females than males (**Figs.1 & 2**) ^45, 46^, we evaluated the roles of the aPVT projection to the mNAcSh in encoding the valence and salience of environmental stimuli. We first asked whether the activity of this circuit was sufficient itself to produce appetitive or aversive responses by assessing voluntary self-stimulation (**Fig. 3**). Male and female mice with an excitatory Gq-DREADD (eDREADD) or empty control vector (CON) in the aPVT-mNAcSh pathway (**Fig. 3a,b; Fig. S2**) performed a home cage preference assay in which they had access to one bottle of water and one bottle of the eDREADD ligand clozapine N-oxide (CNO) dissolved in water (**Fig. 3c**). eDREADD mice had a higher consumption of and preference for CNO than CON mice across CNO concentrations, an effect more robust in females than males (**Fig. 3d,e; Fig. S3**). Notably, this phenotype was present from the first test session and consistent across daily sessions (**Fig. S3a,b,e,f**). These data demonstrate that mice readily and persistently self-activate this circuit and suggest that it is a positively reinforcing circuit, particularly in females. We also assessed the behavioral role of the aPVT-mNAcSh circuit using a sucrose preference test (**Fig. 3f-h; Fig. S4; Fig. S5a,b**). eDREADD, but not CON, mice had higher sucrose consumption and preference following CNO (3 mg/kg) administration than saline vehicle administration for 0.5% sucrose (**Fig. 3g,h; Fig. S4**), and again this phenotype was more robust in females than males. eDREADD mice also had increased preference for 1% sucrose following CNO (**Fig. S5a,b**), confirming that activation of the aPVT-mNAcSh circuit promotes appetitive behavioral responses to rewarding stimuli. We also tested whether the activation of this circuit modulates behavioral responses to negatively valenced stimuli using a Pavlovian fear conditioning paradigm (**Fig. 3i**). eDREADD females, but not males, displayed a blunted fear response (freezing) compared to CON mice during tone presentations during acquisition on day 1 in the presence of CNO, as well as during context retrieval on day 2 in the absence of CNO (**Fig. 3j,k**). eDREADD activation did not affect pain sensitivity (**Fig. S5c,d**), suggesting that the blunted fear response in females may be due to a reduction in the aversiveness of the foot shock while the eDREADD was activated during conditioning rather than alteration in pain perception.

**Fig. 3:**
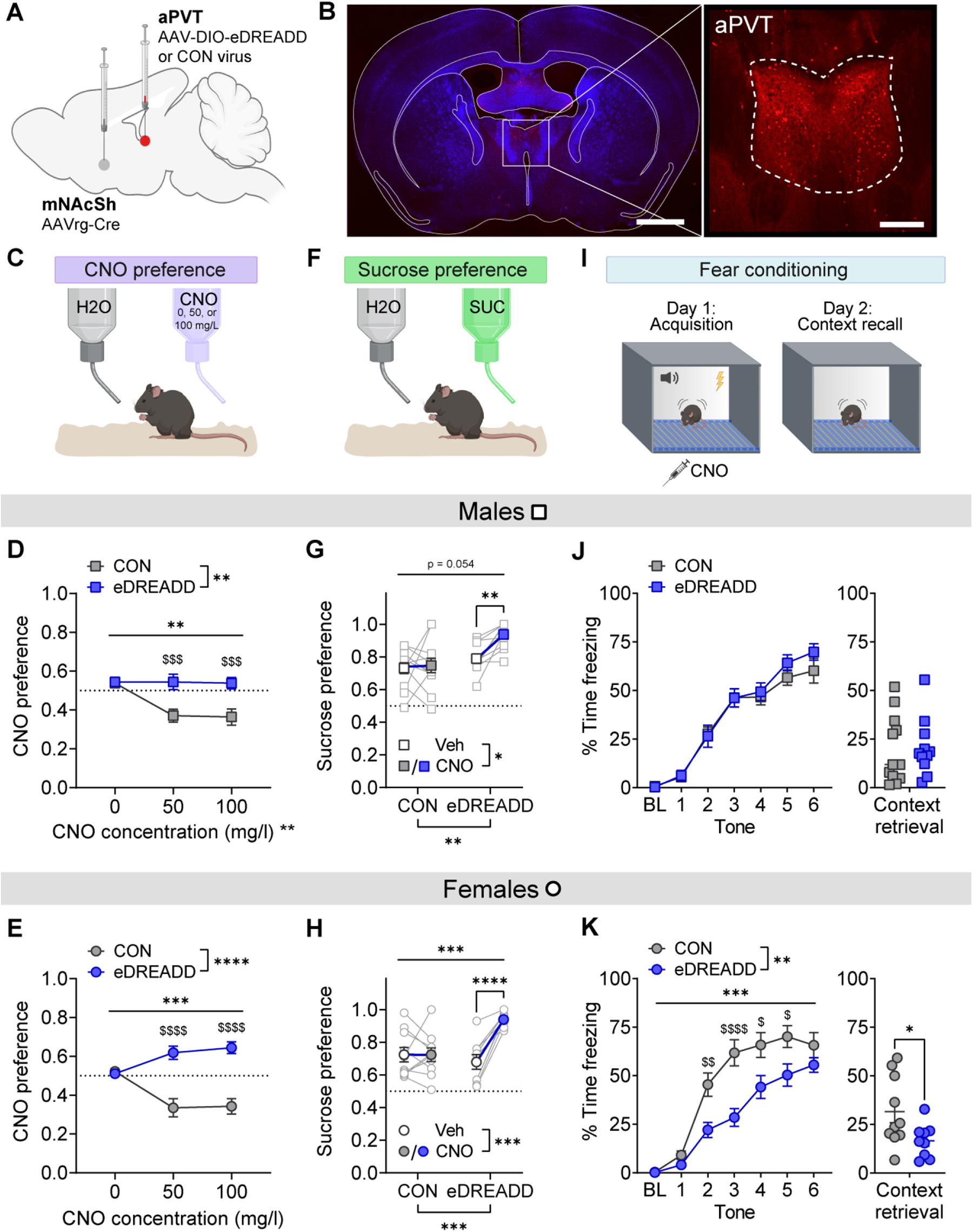
Activity of the aPVT-mNAcSh circuit is positively reinforcing in both sexes, particularly in females. **a)** Schematic of the intersectional viral strategy to express an excitatory Gq-DREADD (eDREADD) or control vector (CON) in the aPVT-mNAcSh circuit. **b)** Representative images of eDREADD expression in the aPVT. Scales bars = 2,500 µm (left) and 500 µm (right) **c-e)** Clozapine N-oxide (CNO) preference test (illustrated in **c**), in which preference for consumption of the eDREADD ligand CNO dissolved in water compared to a water (CNO solution intake / (CNO solution + water intake)) is measured across increasing concentrations of CNO. Male (**d**) and female (**e**) eDREADD mice displayed greater CNO preference compared to CON mice. **f-h)** Sucrose preference test (illustrated in **f**), measured by preference for 0.5% sucrose solution over water (0.5% sucrose intake / (0.5% sucrose + water intake)). Male (**g**) and female (**h**) eDREADD mice, but not CON mice, displayed an increase in sucrose preference following CNO (3 mg/kg, i.p.) compared to saline vehicle (Veh) administration. **i-k)** Fear conditioning assay (illustrated in **i**), with % time freezing during tone presentations preceding foot shock across fear conditioning (blocks of two tones) in the presence of CNO on day 1 (left) and freezing in the fear-conditioned context during retrieval on day 2 (right). eDREADD activation did not alter fear responses in males (**j**) but impaired fear responses during conditioning and context retrieval in females (**k**). \**P* < 0.05, \*\**P* < 0.01, \*\*\**P* < 0.001, \*\*\*\**P* < 0.0001 for ANOVA main effects, interactions, post hoc paired t-tests between Veh and CNO, and unpaired t-tests between CON and eDREADD mice during context retrieval as indicated; ^$^*P* < 0.05, ^$$^*P* < 0.01, ^$$$^*P* < 0.001, ^$$$$^*P* < 0.0001 in post hoc unpaired t-tests between CON and eDREADD mice during fear conditioning.

### aPVT presynaptic terminals in the mNAcSh are tuned to encode valance

These results indicate that the aPVT-mNAcSh pathway is a positively reinforcing circuit that can be more robustly activated to drive positive valence in females (**Fig. 3**), an effect that may be related to our finding that glutamatergic synaptic transmission is more robust in females (**Fig. 2**), and particularly interesting given that females have been largely excluded from previous studies. The literature shows that the PVT plays a role in behavioral responses following conditioning to positively and negatively valenced stimuli, but little is known about its roles in innate, unlearned behavioral responses to stimuli^29, 35, 40, 47, 48^. Therefore, to understand how the aPVT sends valence information to the mNAcSh, we monitored the calcium activity of aPVT synaptic terminals in the mNAcSh during odorant stimulus presentations using GCaMP7c fiber photometry (**Fig. 4a-c**). Female mice were presented with water (neutral stimulus, N) followed by peanut butter (PB, positively valenced stimulus, +) or the fox predator odor trimethylthiazoline (TMT, negatively valenced stimulus, -) for three sec once every 30 sec. Initial presentations of both PB and TMT, but not water, elicited a robust aPVT terminal GCaMP response (**Fig. 4d-f**), consistent with the literature suggesting that the PVT is acutely activated by salient stimuli of both + and – valence^34, 49, 50^. Notably, calcium activity responses were aligned to odorant stimulus presentation onset and remained elevated for many seconds after removal of the odorant stimulus. GCaMP responses were rapidly scaled down across subsequent presentations of TMT but remained large across PB presentations; thus, the steady-state GCaMP response in aPVT terminals differentiated the + and – odorants, with large responses to + and minimal responses to – stimuli (**Fig. 4d-f; Fig. S6**). These fiber photometry results are consistent with our chemogenetic manipulation results showing that enhancement of the activity state of the aPVT-mNAcSh pathway promoted or induced stimulus preference and chemogenetic inhibition ablated preference and induced aversion. Altogether, they suggest that the level activity of the aPVT-mNAcSh circuit elicited by salient stimuli is essential for valence encoding, perhaps via a threshold-dependent glutamate release mechanism in the mNAcSh.

**Fig 4:**
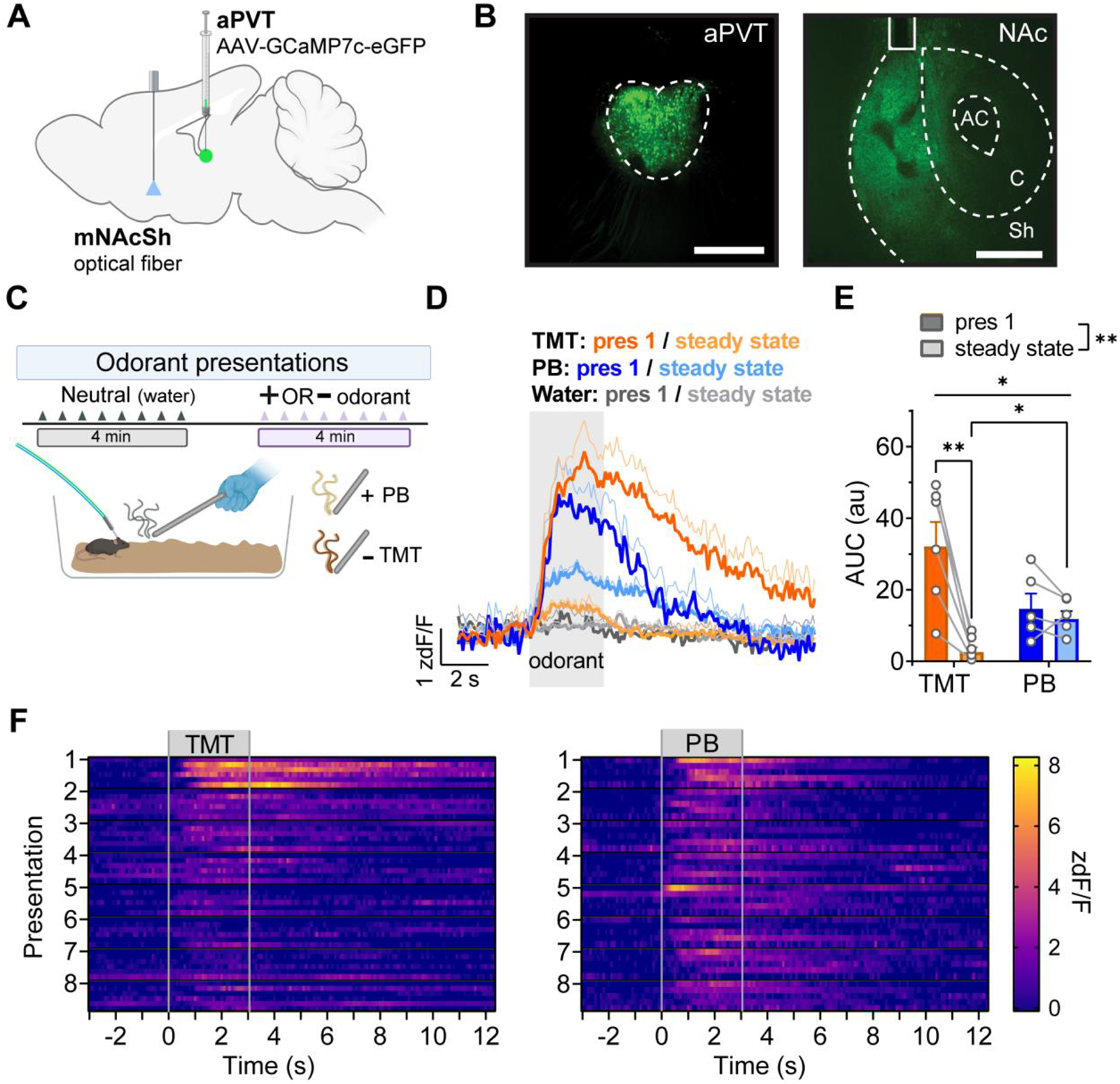
aPVT presynaptic terminals in the mNAcSh are tuned to encode valence. **a)** Schematic of the viral strategy to express the calcium sensor GCaMP7c in the aPVT and implant an optical fiber over the dorsal mNAcSh in female mice. **b)** Representative images of GCaMP expression in the aPVT and mNAcSh, with optical fiber placement over the dorsal mNAcSh. Scale bars = 250 µm and 500 µm, respectively. **c)** Depiction of odorant stimulus presentation paradigm. **d)** GCaMP activity in aPVT synaptic terminals in the mNAcSh time-locked to odorant stimulus presentations, where steady state is represented by the average signal magnitude across presentations 5-8. **e)** Area under the curve for time-locked GCaMP signals shown in **d**. **f)** Heatmaps for GCaMP responses to TMT and PB across stimulus presentations 1-8. Each subrow within presentation represents an individual mouse. \**P* < 0.05, \*\**P* < 0.01 for 2xRM mixed model main effects, interactions, and post hoc paired t-tests as indicated.

### Functional innervation of pPVT neurons to mNAcSh subregions

While our anatomical tracing data suggests that the aPVT more robustly projects to the mNAcSh compared to the pPVT, prior studies in male rodents have shown that the pPVT and its projection to the NAc participates in the regulation of many behaviors, especially stress-sensitive behavioral responses^51–53^. To begin to assess the pPVT-mNAcSh function in both sexes, we used the same optogenetic + slice electrophysiology approach as for the aPVT (**Fig. 2**) to evaluate the strength of pPVT-mNAcSh synapses in the dorsal and ventral subregions in males and females (**Fig. 5a,b**). We found that the amplitudes of oEPSCs in mNAcSh neurons in response to optically-evoked glutamate release from axon terminals of ChR2+ pPVT neurons was robust in both sexes and higher in ventral than in dorsal mNAcSh neurons, particularly in females (**Fig. 5c**). When we directly compared oEPSC magnitude in dorsal and ventral mNAcSh neurons within the same slice using a relative ratio of the dorsal to ventral oEPSC amplitude, we found that this ratio was significantly lower in females compared to males and below 0.50 in females (**Fig. 5d**). These data are consistent with previous studies showing that the pPVT preferentially innervates the ventral mNAcSh^53, 54^. The PPR of oEPSCs from the pPVT input was similar between males and females (**Fig. 5e**), in contrast to what we observed for the aPVT input (**Fig. 2e**), and across the dorsal and ventral mNAcSh, suggesting that the probability of synaptic glutamate release from pPVT terminals is consistent across mNAcSh subregion and sex.

**Fig. 5:**
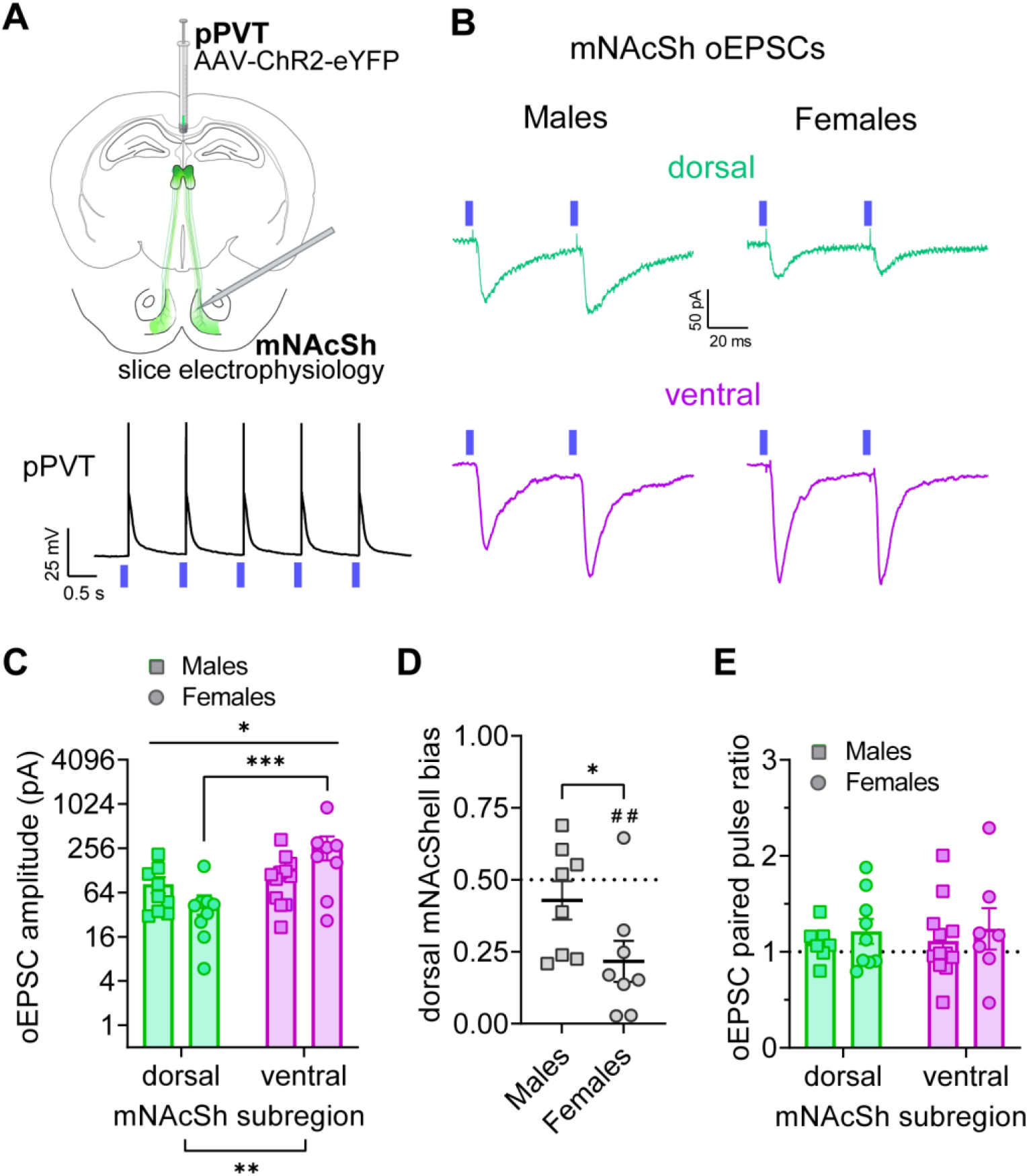
pPVT neurons display more robust excitatory synapses with ventral mNAcSh than dorsal mNAcSh neurons. **a)** Top: Schematic of strategy to express ChR2 in the pPVT-mNAcSh circuit and perform *ex vivo* slice electrophysiology recordings in the mNAcSh. Bottom: Representative traces of 470 nm LED optically-elicited action potentials pPVT cell bodies during current-clamp recordings used to functionally confirm ChR2 expression in pPVT neurons. **b)** Representative traces of optically-evoked excitatory postsynaptic currents (oEPSCs) in neurons in the dorsal (green) and ventral (purple) mNAcSh in response to ChR2-elicited glutamate release from aPVT presynaptic terminals in males (left) and females (right). **c)** Amplitude of oEPSCs in ventral mNAcSh neurons than dorsal mNAcSh neurons, especially in females. **d)** Bias in oEPSC amplitude toward the dorsal mNAcSh, measured by the relative ratio of oEPSC amplitude in the dorsal mNAcSh compared to ventral mNAcSh within individual *ex vivo* slices (amplitude^dorsal^ / (amplitude^dorsal^ + amplitude^ventral^). This ratio is significantly below 0.50 in females. **e)** The paired pulse ratio (PPR) of oEPSCs in dorsal and ventral mNAcSh neurons is similar between sexes and near 1.0. \**P* < 0.05, \*\**P* < 0.01, \*\*\**P* < 0.001 for ANOVA main effects and interaction and post hoc t-tests as indicated; ^##^*P* < 0.01 for one-sample t-tests compared to 1.0 within each group.

### The pPVT-ventral mNAcSh circuit is a salience indicator

We next evaluated the behavioral role of the pPVT-mNAcSh circuit using an intersectional chemogenetic strategy (**Fig. 6a,b; Fig. S8**). Mice with an excitatory Gq-DREADD (eDREADD) or empty control vector (CON) in the pPVT-mNAcSh circuit performed the home cage CNO preference test (**Fig. 6c**). Male, but not female, eDREADD mice had a higher consumption of and preference for CNO than CON mice across CNO concentrations (**Fig. 6d; Fig. S9**). eDREADD activation also enhanced fear responses in males without altering pain sensitivity, but it had no effect on sucrose preference (**Fig. S10a-d;** however, baseline sucrose preference was high and may have occluded an eDREADD effect). Surprisingly, there were no effects of eDREADD activation in females (**Fig. S10e-h**). Together, these data show that males self-stimulate activation of pPVT-mNAcSh neurons and this promotes behavioral responses to aversive stimuli, pointing to a role of the pPVT in enhancing salience in males via its projection to the mNAcSh. To further evaluate the role of the pPVT-mNAcSh in valence and salience in males, we assessed whether chemogenetic activation of the pPVT-mNAcSh circuit could alter appetitive investigation or avoidance of odorant stimuli with innately positive or negative valence (**Fig. 6e**). We found that following CNO administration, CON males investigated female urine (+) more, and TMT (-) less, than water (N). Compared to CONs, eDREADD mice displayed enhanced investigation of female urine (+) and diminished investigation of TMT (-), suggesting that pPVT-mNAcSh pathway activation enhanced the salience of both positively and negatively valenced stimuli to promote innate behavioral responses to both positively and negatively valenced stimuli (**Fig. 6f**). We further probed this pro-salience role using a 0.5% sucrose preference test, finding that administration of either CNO or another eDREADD ligand, Compound 21 (C21), enhanced sucrose preference compared to vehicle in eDREADD but not CON males (**Fig. 6g,h**). Additionally, eDREADD activation with CNO during tone-shock pairings during fear conditioning with low available context information produced an enhancement of the fear response to the context during retrieval (in the absence of CNO) one day later (**Fig. 6i,j**). Altogether, these results provide converging evidence that increasing the activation of mNAcSh-projecting pPVT neurons is sufficient in males, but not females, to provide a salience signal to the mNAcSh that enhances behavioral responses to both positively and negatively valenced stimuli. We followed these behavioral experiments with a circuit and activity mapping approach to understand the brain regions engaged by pPVT-mNAcSh neuron activation (**Fig. S11**). CNO activation of the circuit elicited a robust activation of the pPVT and ventral mNAcSh, but not aPVT or dorsal mNAcSh (**Fig. S11a-c,f**), consistent with the DREADD virus expression (**Fig. S8**). Intriguingly, while mNAcSh-projecting pPVT neurons displayed very little terminal expression in other known projection targets of the pPVT including the central amygdala (CeA), basolateral amygdala (BLA), and medial prefrontal cortex (mPFC), CNO activation of the pPVT-mNAcSh circuit elicited robust Fos expression in the mPFC (**Fig. S11d-f**). These data suggest that the pPVT-mNAcSh circuit’s role in salience may involve indirect circuit engagement of the mPFC.

**Fig. 6:**
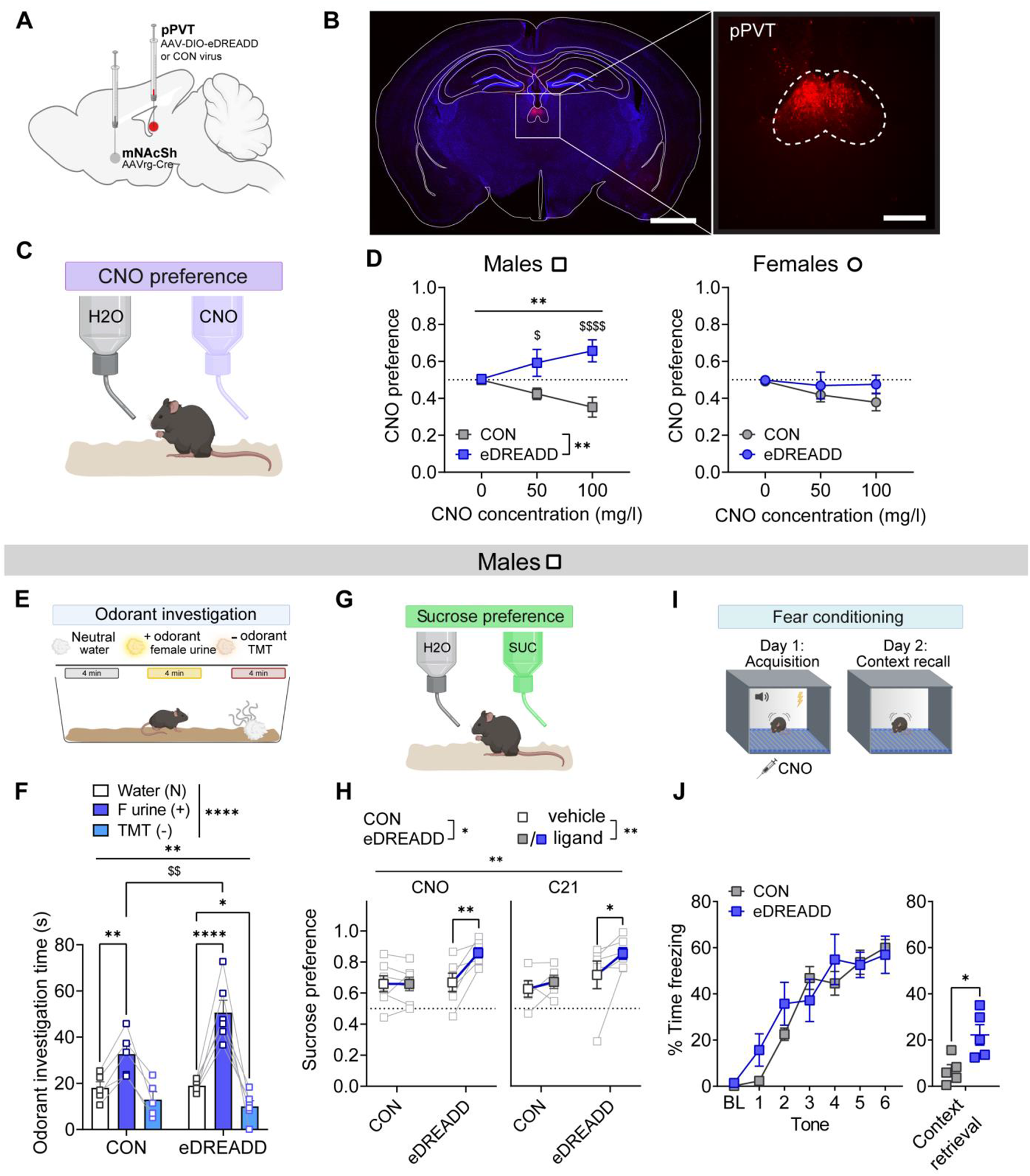
pPVT-ventral mNAcSh circuit activation increases the salience of both positively and negatively valenced stimuli in males. **a)** Schematic of the intersectional viral strategy to express an excitatory Gq-DREADD (eDREADD) or control vector (CON) in the pPVT-mNAcSh circuit. **b)** Representative image of eDREADD expression in the pPVT. Scale bars = 2,500 µm (left) and 500 µm (right). **c-d)** Clozapine N-oxide (CNO) preference test. **c)** Assay illustration. **d)** Male, but not, female eDREADD mice displayed greater CNO preference compared to CON mice across increasing CNO concentrations. **e-j)** Innate and learned behavioral responses to positively and negatively valenced stimuli following CNO (3 mg/kg, i.p.) administration in male CON and eDREADD mice. **e-f)** Odorant investigation assay, with illustration of assay in **e** and data in **f**. Compared to CON males, eDREADD males displayed enhanced investigation of female urine (+) and enhanced avoidance of TMT (-) relative to water (N). **g-h)** Sucrose preference test, with assay illustration in **g** and data in **h**. eDREADD mice, but not CON mice, displayed an increase in 0.5% sucrose preference following CNO or Compound 21 (C21, alternative eDREADD agonist) administration compared to vehicle (Veh) administration. **i-j)** Fear responses during conditioning and context retrieval in a low context fear paradigm, with assay illustration in **i** and data in **j**. % Time freezing to tones (blocks of two tones) during acquisition on day 1 following CNO administration were not altered by eDREADD activation (left), but freezing was higher in eDREADD mice the following day upon exposure to the context in the absence of CNO (right). \**P* < 0.05, \*\**P* < 0.01, \*\*\*\**P* < 0.0001 for ANOVA main effects, interactions, and post hoc paired and unpaired t-tests as indicated. ^$^*P* < 0.05, ^$$^*P* < 0.01, ^$$$$^*P* < 0.0001 in post hoc unpaired t-tests as indicated.

To determine whether the pPVT-mNAcSh circuit is necessary for salience encoding, we performed an intersectional approach to express an inhibitory hM4Di DREADD (iDREADD) in the pathway in males and females (**Fig. 7a,b; Fig. S12**). Following CNO administration, CON males and females investigated PB (+) more, and TMT (-) less, than water (N). In contrast, iDREADD mice displayed no preference for PB (+) and had an overall decreased investigation of odorant stimuli compared to CONs in both males (**Fig. 7c; Fig. S13a,b**) and females (**Fig. 7d; Fig. S13c,d**). These data suggest that stimulus salience, and thus motivated exploration, was decreased when pPVT-mNAcSh circuit activity was lower. In a sucrose preference test, iDREADD activation with CNO decreased sucrose preference in both sexes without an effect of CNO in CON mice (**Fig. 7e,f**). iDREADD activation during fear conditioning with high available context information produced a decrease in the fear response to the context during retrieval (in the absence of CNO) one day after conditioning in both sexes (**Fig. 7g,h**). Altogether, these results provide converging evidence that activity of the pPVT-mNAcSh circuit is a necessary salience indicator in both males and females that guides both innately-programmed and learned behavioral responses to stimuli regardless of valence.

**Fig. 7:**
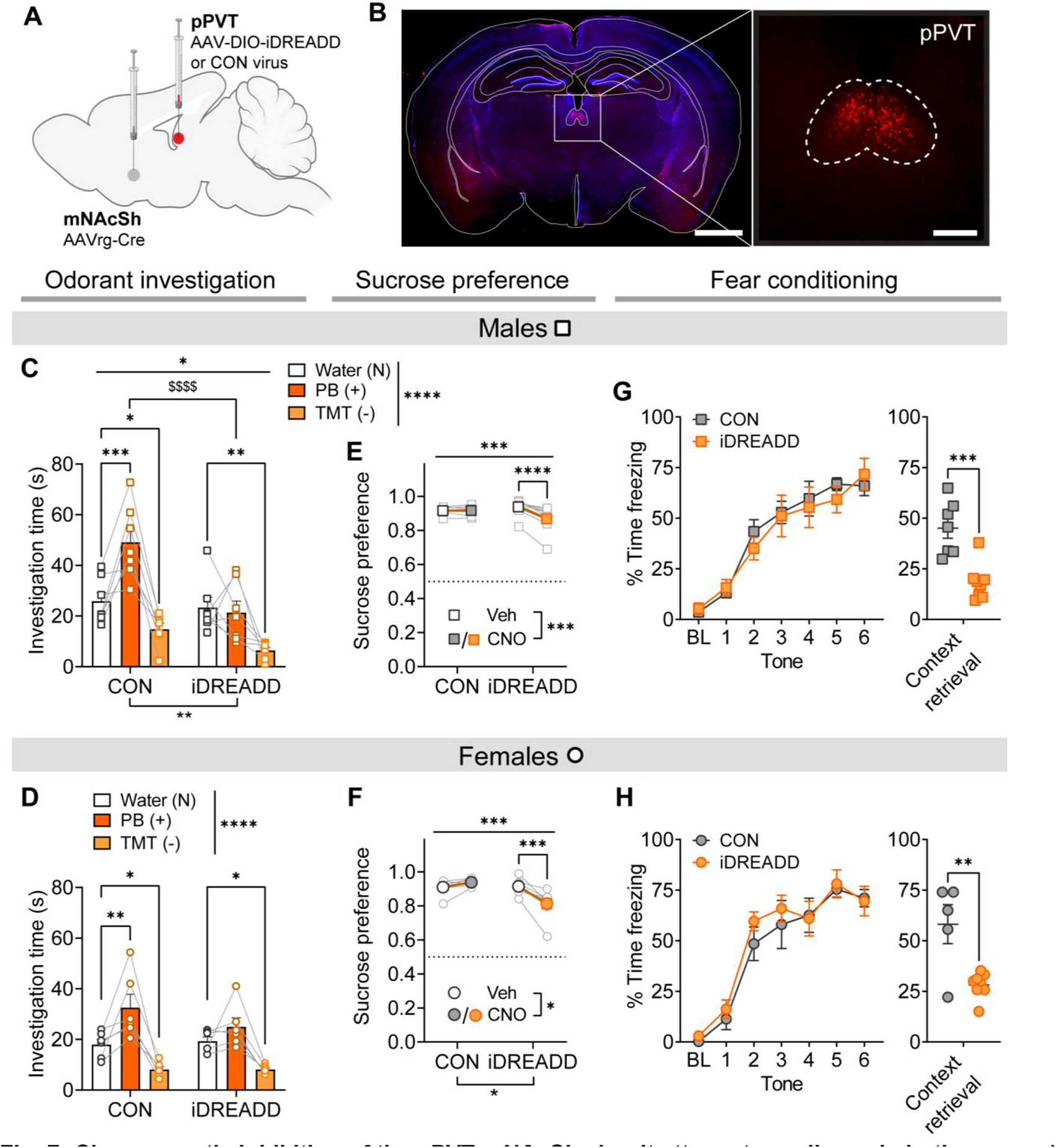
Chemogenetic inhibition of the pPVT-mNAcSh circuit attenuates salience in both sexes. **a)** Schematic of the intersectional viral strategy to express an inhibitory hM4D-Gi DREADD (iDREADD) or control vector (CON) in the pPVT-mNAcSh circuit. **b)** Representative image of iDREADD expression in the pPVT. Scale bars = 2,500 µm (left) and 500 µm (right). **c-d)** Odorant investigation assay following CNO (3 mg/kg, i.p.) administration in males (**c**) and females (**d**). Compared to CON mice, iDREADD mice displayed an overall reduction in odorant investigation time and a loss of preference for peanut butter (PB, +) over water. **e-f)** Sucrose preference test in males (**e**) and females (**f**). iDREADD mice, but not CON mice, displayed a suppression in 1% sucrose preference following CNO compared to vehicle (Veh) administration. **g-h)** Fear responses during conditioning and context retrieval in males (**g**) and females (**h**). % time freezing to tones (blocks of two tones) during acquisition on day 1 following CNO administration (left) was unchanged but % time freezing to context during retrieval on day 2 in the absence of CNO (right) was decreased in iDREADD mice. \**P* < 0.05, \*\**P* < 0.01, \*\*\**P* < 0.001, \*\*\*\**P* < 0.0001 for ANOVA main effects, interactions, and post hoc t-tests as indicated; ^$$$$^*P* < 0.0001 in post hoc t-test between CON and iDREADD mice.

## Discussion

The PVT projects to many brain regions, with the mNAcSh being a primary target. Previous studies have demonstrated various roles of the PVT-NAc circuit, primarily in male rodents, in conditioned behaviors related to seeking and consumption of natural and drug rewards, as well as adaptive fear responses such as active avoidance^5, 6, 29, 40, 55^. Parallel, robust literatures have separately examined the specific roles of anatomical subregions of the PVT^34, 35, 41^. Here, we overlay the intra-PVT region organization (aPVT vs. pPVT) onto the anatomy and function of PVT-mNAcSh circuits to understand the relationship between these components and behavioral control. We describe parallel excitatory projections from the aPVT and pPVT that provide key, but different, types of information about external stimuli to the nucleus accumbens to shape adaptive behavioral responses. We found that the aPVT and pPVT, through their glutamatergic synaptic inputs to the mNAcSh, convey information about the valence and salience of stimuli, respectively. Importantly, the roles of these thalamo-accumbal pathways in stimulus feature encoding are essential for adaptive behavioral responses to stimuli with innately programmed salience and valence, such as odors of opposite sex conspecifics (positively valenced) and predators (negatively valenced), and extend to stimuli learned to be salient and have valence, such as the inherently neutral context that becomes a salient and aversive stimulus following its pairing with foot shock during fear conditioning. These results add important refinement about the topographical organization and temporal dynamics for the role of the PVT-NAc circuit previously shown to a play a role in behavioral responses following conditioning.

We found that the activity of the aPVT-mNAcSh circuit plays a role in positive valence encoding (**Fig. 3**) and was positively reinforcing *per se*, as mice self-stimulated the circuit via voluntarily consumption of the excitatory DREADD ligand CNO (**Fig. 3c-e; Fig. S3**). The role of this circuit in valence encoding that extends from innately programmed behavioral responses to more complex learned behaviors and across modalities (**Figs. 3 & 4**) suggests that its set point in activity is adaptively programmed to be responsive to the environment. Consistent with this, we found that while the aPVT-mNAcSh circuit responds robustly to both positive and negative stimuli, it conveys information about the valence of stimuli through rapid tuning of presynaptic terminal activity to produce larger steady state responses to positively valenced than negatively valenced stimuli in the mNAcSh (**Fig. 4; Fig. S6**). This may be related to the known complex role of the topographical organization of the mNAcSh in aspects of motivated behavior. For example, a portion of the anterior mNAcSh is known to be a hedonic hotspot, such that its activity increases “liking”, while the posterior mNAcSh a hedonic coldspot^1–3, 42, 56^, effects that are dependent on glutamatergic synaptic transmission^2, 3, 42, 56–58^. Here, we primarily targeted the anterior half of the mNAcSh, suggesting that the aPVT may be an important source of glutamate for valence encoding in the anterior mNAcSh. Whether aPVT input to the more posterior aspects of the mNAcSh would differentially affect behavior remains unknown. Additionally, the roles of subpopulations of mNAcSh neurons and opioid receptor signaling in reward and aversion depend on anterior-posterior and dorsal-ventral location within the NAcSh^59, 60^. For example, dynorphin (putative D1 receptor+) neuron activity is positively reinforcing in the dorsal mNAcSh and aversive in the ventral mNAcSh^60^. There may be aPVT-Dyn neuron interactions playing a role in valence encoding, as dynorphin/D1R+ neurons are postsynaptic targets of the PVT^27, 38^ and locally released dynorphin may act upon aPVT synaptic terminals. As our electrophysiology data show that female aPVT-mNAcSh neurons have a higher probability of synaptic glutamate release than males (**Fig. 2**), and our behavioral data show that chemogenetic circuit activation elicits a more robust behavioral modulation in females than males (**Fig. 3**), gating of aPVT glutamate release in the NAc may be one mechanism for sex differences in valence encoding.

We also found that the role of the aPVT-mNAcSh in valence encoding may extend to fear responses in females (**Fig. 3k**), particularly interesting given the emerging literature showing that the NAc is an important modulatory hub for fear behavior. For example, the activity of cholinergic interneurons in the NAc is required for fear responses during context recall^61^, while dopamine signaling at D1 receptors inhibits this behavior^62^. Importantly, glutamatergic synaptic transmission in the mNAcSh is required for adaptive fear responses^1–3^, and the NAc is involved in the rapid scaling of fear responses to the current level of threat^63, 64^. We found here that the activity aPVT glutamatergic synaptic inputs to the NAc is rapidly tuned to match and encode similar stimulus properties and these may be important for fear responses in females. In contrast, we found that the pPVT-mNAcSh circuit enhanced fear responses in a manner consistent with its role in salience encoding, as its activation status was causally related to behavioral expression in both positive (+ odorants, sucrose preference) and negative (-odorants, fear) experimental settings. Interestingly, the endogenous activity of the pPVT-mNAcSh circuit during behavior was necessary for salience encoding in both sexes, but activation of the circuit above its endogenous activity level was only sufficient to enhance stimulus salience in males (**Figs. 6 & 7**). Further, males, but not females, preferentially consumed CNO to activate the pPVT-mNAcSh pathway (without engaging the aPVT valence circuit; **Fig. 6d**), even though the anatomical and functional density of the projection was robust in both sexes (**Figs. 1 & 5**). This suggests that this projection is highly engaged during behavior and that the tone of activity in the pPVT-mNAcSh circuit may be higher during behavior in females than males. These results have implications for sex-dependent behavioral output depending on the inherent and learned salience of animals’ current environmental surroundings. As we also found that the aPVT projection to the mNAcSh, at least in females, inhibits fear responses, our data point to independent and perhaps opposing functional roles of the aPVT and pPVT projections to the NAc in shaping fear behavior.

Our finding that the activity of the pPVT-mNAcSh circuit during fear conditioning is important for an adaptive fear response (freezing) during context recall the following day (**Figs. 6 and 7**) is similar to a previous study examining the pPVT projection to the central amygdala, another primary target of pPVT neurons^29^. This is a particularly interesting similarity given that we find that the ventral mNAcSh-projecting population does not significantly innervate the CeA (via bifurcating axons), nor its activation significantly recruit the CeA. Overall, pPVT neurons respond to footshock and predictive cues^6, 34, 35^. Given the emerging role of the NAc in fear behavior and density of pPVT projections to it, it may be the case that the important role of the pPVT in fear behavior is through separate projection populations conveying different types of relevant information to shape behavioral output. Interestingly, a subpopulation of pPVT-mNAcSh neurons expressing dopamine D2 receptors (D2Rs) has recently been shown to play a role in the expression of conditioned active avoidance of a footshock-paired context; terminal GCaMP activity was increased when mice performed the adaptive response (escape) but inhibited when they performed the less adaptive behavioral response (freeze)^35^. Together these data speak to the importance of the pPVT-mNAcSh circuit in appropriate behavioral responses in a fearful setting. Future studies are needed to determine whether the overall pPVT-mNAcSh and D2R+ subpopulation play similar roles across unconditioned (as in our study) and conditioned responses.

While previous studies have generally focused on the roles of PVT subpopulations in conditioned behaviors, we found that the aPVT and pPVT each, through their mNAcSh projections, play a role in the encoding of stimulus features that affect unconditioned and conditioned responses. A few other studies provide insight regarding the responses of PVT subpopulations to stimuli in unconditioned or early learning time points. One study found that GCaMP responses in PVT-NAc neurons in males were, on average, decreased during sucrose consumption early in operant conditioning; however, there was heterogeneity across the population, with some neurons displaying increased activity^40^. This spectrum of activation may depend on anatomical location within the PVT. For example, recent studies showed that DRD2+ neurons in mice, primarily concentrated in the pPVT, were acutely activated by aversive stimuli but inhibited by positively valenced stimuli ^34, 35^, suggesting they are valence sensitive; in contrast, DRD2-neurons, more concentrated in the aPVT and with a different circuit organization pattern than DRD2+, were inhibited by all stimuli, suggesting they are salience-sensitive^34^. These subpopulations of PVT neurons were shown to differentially and reciprocally engage with the IL vs. PL medial PFC. Intriguingly, we found here that pharmacogenetic activation of pPVT-ventral mNAcSh neurons, found to have few collaterals projecting directly to the mPFC, robustly activated both IL and PL subregions of the mPFC without engaging the aPVT (**Fig. S11**). This suggests that the pPVT-ventral mNAcSh circuit under investigation here may indirectly engage the reciprocal cortico-thalamic loops, with implications for behavioral regulation^31, 34, 65, 66^ .

In contrast to what has been shown for these molecularly distinct populations in male/sex-unspecified mice, an examination of aPVT vs. pPVT neurons in male rats showed that aPVT neuron GCaMP activity is increased upon presentation of cues predicting sucrose availability or footshock, as well as footshock itself (but not sucrose consumption)^6, 67^. However, aPVT GCaMP activity during conditioned stimulus presentation was positively correlated with conditioned responding across learning for sucrose but not fear. In contrast, pPVT neurons responded robustly to both the positively and negatively valenced stimuli and cues predicting them, and its activity during conditioned stimulus presentation was positively correlated with conditioned responding across learning for both sucrose and fear tasks. Together, these data suggest that the aPVT may serve as a valence indicator, while the pPVT may be a salience indicator, similar in many ways to what we observed here for the roles of PVT-mNAcSh circuits.

While the specific responses of aPVT and pPVT neuron populations to positively and negatively valenced stimuli varies across studies, together their results a suggest a complex functional interaction between the aPVT and pPVT in encoding the salience and valence of stimuli that is critical to the regulation of adaptive motivated behaviors. These interactions may lie in their convergence upon common targets, including the mNAcSh, with behavioral roles occurring in anatomically defined and sex-dependent ways. In our studies, we found that the aPVT and pPVT both provide synaptic input to the dorsal and ventral subregions of the mNAcSh. However, there were different sex-dependent and independent characteristics of aPVT and pPVT inputs, such as higher probability of synaptic glutamate release from aPVT (but not pPVT) terminals in females than in males, and a bias toward ventral mNAcSh for pPVT (but not aPVT) inputs (**Figs. 2 & 5**). Future studies will be needed to understand how the aPVT and pPVT inputs are integrated within and across dorsal and ventral mNAcSh subregions, as well as across the anterior-posterior extent of the mNAcSh, to shape behavioral output in males and females. In addition, future consideration should be given to understanding how broader topographical mapping of PVT subregions described in males^45, 46^ is organized in females.

## Methods

### Subjects

Male and female C57BL/6J mice were purchased from Jackson Laboratories (stock # 000664; Bar Harbor, ME, USA) at eight weeks of age and housed under a reverse circadian 12 h:12 h light:dark cycle with lights off at 7:30 am and *ad libitum* access to food and water. Mice were singly housed at least one week prior to behavioral testing and remained singly housed throughout the experimental procedures. Experiments were conducted during the dark phase of the light:dark cycle. All procedures were conducted with approval of the Institutional Animal Care and Use Committee at Weill Cornell Medicine following the guidelines of the NIH Guide for the Care and Use of Laboratory Animals.

### Stereotaxic surgeries

Mice underwent stereotaxic surgery for viral injection and fiber implantation as previously described^24^. Mice were anesthetized with 2% isoflurane (VetEquip, Livermore, CA) in 0.8% oxygen in an induction chamber (VetEquip, Livermore, CA) and placed in an Angle Two mouse stereotaxic frame (Leica Biosystems, Wetzlar, Germany). Mice were given a subcutaneous injection of meloxicam (2 mg/kg) for preemptive analgesia and 0.1 mL of 0.25% Marcaine around the incision site. To selectively express viral constructs, we infused through a Neuros 7000 series 1 µL Hamilton syringe with 33-gauge needle (Reno, NV) connected to a remote automated microinfusion pump (KD Scientific, Holliston, MA) at a rate of 25-50 nL/min, using the following stereotactic coordinates: aPVT (A/P: -0.46mm, M/L: 0.00mm, D/V: -3.35mm, 300nL for all viruses except 750 nL for GCaMP7c), pPVT (A/P: -2.14mm, M/L: 0.00mm, D/V: -2.82mm, 200nL), mNAcSh (A/P: 1.30mm, M/L: 0.50mm, D/V: -4.25mm (Figs. 1 & 2) or -4.75mm (Figs. 6 & 7), 250-300nL each). Following infusion, the needle was left in place for 10 min and then slowly manually retracted to allow for diffusion and prevent backflow of virus. For fiber photometry experiments, mice also received fiber optic cannula (400 μm core, 0.53 NA with angled tip; Doric Lenses) implantation over the mNAcSh, secured to the skull using MetaBond and dental cement. Mice were continuously monitored for at least 30 minutes post-surgery to ensure recovery of normal breathing patterns and checked daily thereafter.

### Viruses

For DREADD experiments, mice received bilateral intra-mNAcSh injections of a retrograde Cre virus (AAVrg-hSyn.HI.EGFP-Cre.WPRE.SV40 or AAVrg-hSyn.Cre.WPRE.hGH, 1.17-2.1×10^13^ gc/mL) and an intra-aPVT or intra-pPVT injection of an eDREADD (AAV2-hSyn-DIO-hM3D(Gq)-mCherry, 6.5×10^12^-2×10^13^ gc/mL), iDREADD (AAV2-hSyn-DIO-hM4D(Gi)-mCherry, 4.9×10^12^ gc/ml), or control virus AAV2-hSyn-DIO-mCherry (4.7×10^12^ gc/mL). For anterograde tracing and slice electrophysiology mice received intra-PVT injections of ChR2 (AAV9-CaMKIIa-hChR2(H134R)-eYFP.WPRE.hGH, 1.5×10^13^ gc/mL). For fiber photometry studies, mice received intra-aPVT injections of GCaMP7c (AAV1-Syn-jGCaMP7c (1.1×10^13^ gc/mL). For retrograde tracing experiments, mice received unilateral injections of retrogradely expressed fluorophore-tagged viruses AAVrg-CAG-GFP (7.0×10^12^-2.2×10^13^ vg/mL) or AAVrg-CAG-tdTomato (1.02-1.6×10^13^ vg/mL).

### Chemogenetic manipulations during behavior

Mice were injected with a retrogradely trafficked Cre-expressing virus in the mNAcSh and a Cre-dependent eDREADD, iDREADD, or empty control vector in the aPVT or pPVT. After three weeks to allow for viral expression, mice underwent behavioral tests to evaluate the role of the PVT-mNAcSh projections in behaviors with positively and negatively valenced stimuli and varying degrees of salience. Mice were habituated to experimenter handling prior to behavioral testing. Consummatory assays were conducted under red light, and all other behavior assays were conducted under 250 lux lighting conditions^68–70^. All behavioral experiments commenced three hours into the dark cycle, and each behavioral apparatus was thoroughly cleaned with 70% ethanol before each trial. For chemogenetic manipulation experiments, mice received vehicle (0.9% sterile saline, i.p.) or DREADD ligand dissolved in vehicle (CNO, 3 mg/kg, i.p., or Compound 21, 17 mg/kg, i.p.) injections 40 min prior to behavioral testing.

#### CNO preference

CNO consumption and preference testing was performed in a similar fashion to previous studies^71^. Home cage water bottles were removed and mice were given access to two bottles, one with tap water and one with CNO dissolved in water, for four hr/day for four consecutive days per week. Water and CNO bottle placement was switched daily to avoid potential innate or learned side preferences. CNO concentration increased each week across three weeks (0, 50, 100 mg/L). CNO and water consumption were calculated based on bottle weight before and after the session, subtracting out estimated loss based on bottle weights from dummy cages and normalized to the mouse’s bodyweight. CNO concentrations were selected to achieve biologically relevant brain concentrations sufficient to activate centrally-expressed DREADDs, and experiments were performed during a time period in which mice typically consume 30-50% of their total daily fluid consumption in a 4h period.

#### Sucrose preference

Sucrose intake and preference was performed in a similar manner to that described for the CNO preference test. Home cage water bottles were removed and mice were given access to two bottles, one with tap water and one with sucrose dissolved in water (1 or 0.5% w/v) for four hr/day for four consecutive days per week, and water and sucrose bottle side was switched daily. Days 1 and 2 of the week served as a baseline period. Mice received injections of vehicle and CNO on days 3 and 4, respectively, 40 min prior to testing.

#### Fear conditioning

Fear conditioning was performed in an operant box with a stainless-steel grid floor within a sound-attenuating chamber (Coulbourn Instruments, Allentown, PA, USA). Mice received a 3 mg/kg i.p. injection of CNO 40 minutes prior to behavioral testing. Mice were placed in the chamber at the beginning of the test, and following a 5 min habituation period received six pairings of a 30 s, 80 dB tone (conditioned stimulus, CS) co-terminating with a 2 s, 0.5 mA foot shock (unconditioned stimulus, US) separated by pseudorandom intertrial interval (31 – 119 s with an average of 75.5 s). Video tracking and FreezeFrame software (Coulbourn Instruments, Allentown, PA, USA) were used to assess freezing behavior during the 28 s tone prior to shock presentation.

#### Thermal paw withdrawal

Pain sensitivity was measured using a hot plate assay. Mice were confined in a Plexiglas chamber on a metal surface set at 52 °C and the latency to a nociceptive response (licking or shaking of hind paws, jumping) was assessed. To prevent tissue damage, mice were removed from the hot plate after a nociceptive response or upon reaching a cut-off time of 30 seconds. Triplicate measurements 15 min apart were obtained and the average latency per animal calculated.

#### Odorant stimulus investigation

Mice were habituated to a clean home cage with a 1 cm x 1 cm cotton nestlet for 20 min prior to being to odorant stimuli with innately programmed preference or avoidance behavioral responses. Nestlets soaked in water (neutral stimulus, N), opposite sex urine or peanut butter (positive, +), and diluted TMT or 2-IB (negative, -) were sequentially placed on one side of the experimental cage for four min, and the mouse was transferred to its home cage for 5 min between odorants while the experimenter cleaned the cage. Investigation of with the stimulus was measured by the number of bouts and total time the mouse spent sniffing each nestlet. A modified version of this assay was used to assess aPVT-dorsal mNAcSh terminal calcium activity during odorant exposure, described below.

### *in vivo* fiber photometry

Mice expressing GCaMP7c in the aPVT and fiber cannulae over the mNAcSh were habituated to handling and tethering of fiber optic patch cables, as well as to being presented with a cotton swab by the experimenter in the home cage, during the week prior to the odorant stimulus presentation experiment. Optical patch cables were photobleached for at least two hr prior to the start of the experiment. At the start of each fiber photometry recording session, the mouse’s optical fiber cannulae were tethered to the optical patch cable connected to the fiber photometry rig. The experimenter presented a cotton swab with water (N) and then either PB (+) or diluted TMT (-) for three sec once every 30 s for four min, with a several min break between the two odorant presentation trains. The following week, the same procedure was repeated with the other odorant. GCaMP fiber photometry data were collected via TDT Synapse software and analyzed using custom MATLAB and RStudio code developed in our lab. Raw F values in the 465 nm (GCaMP) channel and isosbestic control 405 nm channel for each session (N and +/-odorant) were fit with linear or two-term exponential fits, determined by the fit of the 405 nm channel data, and delta F/F_0_ values calculated and z-scored for analysis. Peak responses within the first four sec and area under the curve for the 12 sec of the onset of odorant presentation (time = 0) were calculated using the three sec baseline period preceding odorant presentation. For steady state responses, traces for presentations 5-8 were averaged.

### Fluorescence immunohistochemistry

Following behavior procedures, mice were deeply anesthetized with pentobarbital (100 mg/kg, i.p.) and transcardially perfused with sterile phosphate-buffered saline (PBS) followed by 4% paraformaldehyde (PFA). Brains were extracted, post-fixed overnight in 4% PFA, and then placed in PBS until they were sliced on the coronal plane in 45 μm sections on a VT1000S vibratome (Leica Biosystems) to determine viral expression and fiber placements. eDREADD and CON mice used for Fos and terminal expression quantification were injected with CNO 90 min prior to sacrifice. Brain sections underwent fluorescence immunohistochemistry using standard procedures^24, 72^. Tissue was washed in phosphate buffer saline (PBS) and then incubated in blocking buffer containing 0.2% Triton X and 1% normal donkey serum in PBS for 1 hr at room temperature, followed by blocking solution containing primary antibodies against mCherry (mouse monoclonal 1:500, Takara Bio) and c-Fos (rabbit polyclonal 1:500, Synaptic Systems) overnight at 4°C. The next day, slices were washed in PBS and incubated in respective secondary antibodies against mouse (donkey Alexa Fluor 568, 1:500, Invitrogen) and rabbit (donkey Alexa Fluor 488, 1:500, Jackson Immuno Research) in blocking buffer for one hr at room temperature. Sections were washed, counterstained with DAPI, mounted using Vectashield hard mount antifade mounting medium (Vector Labs), and stored in the dark at 4°C until imaged on a Nikon Eclipse 80i microscope with monochrome sCMOS camera using NIS-Elements (AR 5.30.05, build 1559) software for DAPI (405 nm), c-Fos (488 nm), and mCherry (568 nm). Terminal brain regions were imaged using the same microscope settings across mice, and CellProfiler software (v 4.2.1, Broad Institute, Inc.) was used to quantify the number of DAPI and c-Fos+ cells and the fluorescence intensity of mCherry.

### Anatomical tracing

Mice were injected unilaterally with a retrogradely trafficking fluorophore-tagged viruses in the mNAcSh and then sacrificed three weeks later and their brains extracted, fixed, and sliced as described above. PVT and mNAcSh slices were counterstained with DAPI and mounted onto glass slides. Images of the PVT across Bregma coordinates were acquired on a Zeiss LSM 880 Laser Scanning confocal microscope (Carl Zeiss, Oberkochen, Germany) and quantified in MetaMorph (San Jose, CA) for the number of DAPI+ nuclei and GFP or tdTomato-expressing cells.

### *ex vivo* electrophysiological recordings

Mice that received intra-aPVT or intra-pPVT injections of ChR2 were rapidly decapitated at least 6 weeks later under isoflurane anesthesia and their brains extracted. Brains were blocked on the coronal plane and acute 300 µM coronal slices prepared and incubated in carbogenated solutions with a pH of 7.35 and osmolarity of 305 ^73^. Sections including PVT or NAc were sliced in *N*-Methyl-D-glucamine (NMDG) artificial cerebrospinal fluid (NMDG-aCSF) at RT containing (in mM): 92 NMDG, 2.5 KCl, 1.25 NaH2PO4, 30 NaHCO3, 20 HEPES, 25 glucose, 2 thiourea, 5 Na-ascorbate, 3 Na-pyruvate, 0.5 CaCl2·2H2O, and 10 MgSO4·7H2O. Slices were transferred to NMDG-aCSF at 32 °C for 12-14 min, and then incubated for at least 1 hr at RT in HEPES-aCSF containing (in mM): 92 NaCl, 2.5 KCl, 1.25 NaH2PO4, 30 NaHCO3, 20 HEPES, 25 glucose, 2 thiourea, 5 Na-ascorbate, 3 Na-pyruvate, 2 CaCl2·2H2O, and 2 MgSO4·7H2O. Slices were placed in the recording chamber and perfused at a rate of 2 ml/min with 30 °C normal aCSF containing (in mM): 124 NaCl, 2.5 KCl, 1.25 NaH2PO4, 24 NaHCO3, 12.5 glucose, 5 HEPES, 2 CaCl2·2H2O, and 2 MgSO4·7H2O, for at least 20 min prior to electrophysiological recordings.

Whole-cell patch-clamp recordings of aPVT and pPVT neurons were performed in current-clamp configuration to confirm appropriate ChR2 expression and function prior to recordings in postsynaptic mNAcSh neurons. PVT neurons were patched using a potassium gluconate-based intracellular recording solution containing (in mM): 135 KGluc, 5 NaCl, 2 MgCl_2_-6H2O, 10 HEPES, 0.6 EGTA, 4 Na-ATP and 0.4 Na-GTP (pH 7.3 and 290 mOsm), and 1-ms 470 nm LED stimulation was used to optically stimulate ChR2+ to elicit action potentials at the cell bodies of patched neurons to confirm sufficient PVT ChR2 expression in the target region (aPVT in **Fig. 4A**; pPVT in **Fig. 8A**). The other subregion was also visually and functionally assessed in the same manner to confirm a lack of ChR2. Glutamatergic synaptic transmission between aPVT/pPVT terminals and postsynaptic dorsal/ventral mNAcSh neurons was measured in voltage-clamp configuration at a holding potential of -70 mV and with picrotoxin (25 µM) in the aCSF bath. In postsynaptic mNAcSh neurons patched using a cesium-gluconate-based intracellular recording solution containing (in mM): 117 D-gluconic acid, 20 HEPES, 0.4 EGTA, 5 TEA, 2 MgCl_2_-6H2O, 4 Na-ATP and 0.4 Na-GTP (pH 7.3 and 290 mOsm), two 2-ms pulses of 470 nm LED stimulation with an ISI of 50 ms were used to evoke glutamate release from PVT terminals while recording optically-evoked postsynaptic currents (oPSCs). Within each slice, recordings were collected from cell(s) in both the dorsal and ventral mNAcSh subregions to assess the relative responses between subregions within Bregma coordinate to the same ChR2 stimulation. Signals were acquired using a Multiclamp 700B amplifier (Molecular Devices), digitized, and analyzed via pClamp 10.4 or 11 software (Molecular Devices). Input resistance and access resistance were continuously monitored throughout experiments, and cells in which properties changed by more than 20% were not included in data analysis. oEPSC amplitude for the first and second pulses was calculated in Clampfit 11.

### Statistical analyses

Statistical analyses were performed in GraphPad Prism 9. Data in each group for all dependent measures were checked for their distributions in raw and log space within the group and equality of variance across groups and analyzed accordingly. Synaptic transmission data were normally distributed in log space, and all other data in raw space. Welch’s correction was used when variances between groups were unequal for planned t-tests. Statistical comparisons were always performed with an alpha level of 0.05 and using two-tailed analyses. Two-way and three-way repeated measures ANOVAs (RM-ANOVAs) were used to examine the effects of cycle, time, dose, tone, brain region, and other repeated variables between mice in different experimental conditions; mixed effects models were used when one or more matched data point was unavailable. Post hoc direct comparisons following ANOVAs were performed using unpaired or paired t-tests with Holm-Šidák (H-S) correction for multiple comparisons, and corrected p values are reported. Statistical analyses are reported in **Table S1**. Data in figures are presented as mean ± SEM; raw data points are included in all figures except some repeated measures graphs in which there were too many raw data points to be clearly represented.

## Acknowledgments

This work was supported by NIH/NIAAA grants R00 AA023559 and R01 AA027645 to K.E.P and F31 AA029293 to L.J.Z, a Brain and Behavior Research Foundation NARSAD Young Investigator Award (26608) to K.E.P, and NIH/NIDA grant T32 DA03980 to J.K.R.I. and L.J.Z. We thank Scottie Nelson and John Miller for technical assistance.

## Author contributions

J.K.R.I., P.U.H., and K.E.P. designed the experiments and wrote the manuscript; K.E.P oversaw all studies; all authors conducted experiments, analyzed data, and edited and approved the manuscript.

