## Supplementary Figures for "Valence and salience encoding by parallel circuits from the paraventricular thalamus to the nucleus accumbens"

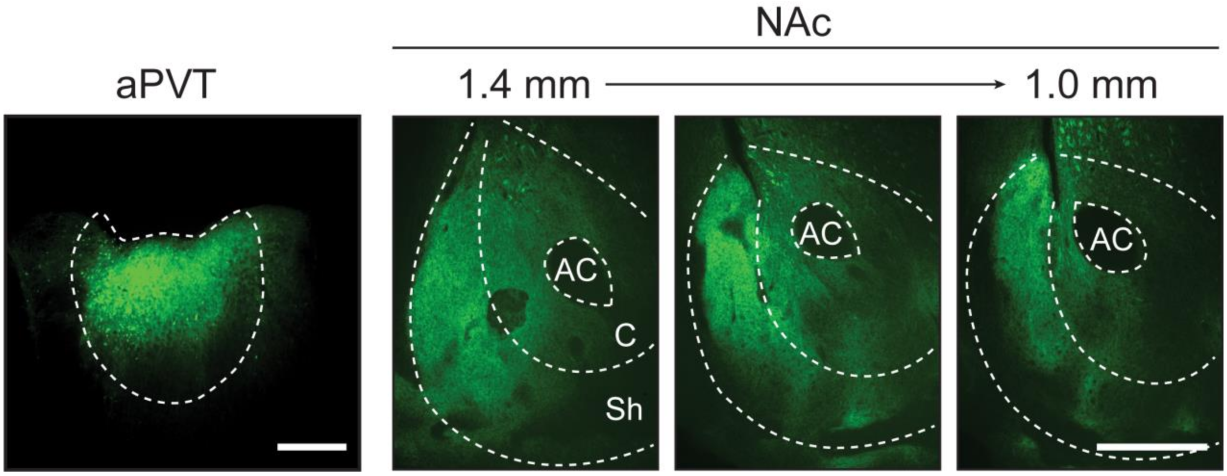


**Supplementary Fig. 1 (related to Fig. 2a): Representative images of ChR2 expression in the aPVT and across the A-P extent of the nucleus accumbens in a female mouse injected with ChR2 in the aPVT.** AC = anterior commissure, C and Sh = core and shell of the nucleus accumbens, respectively. Scale bars = 500 µm.

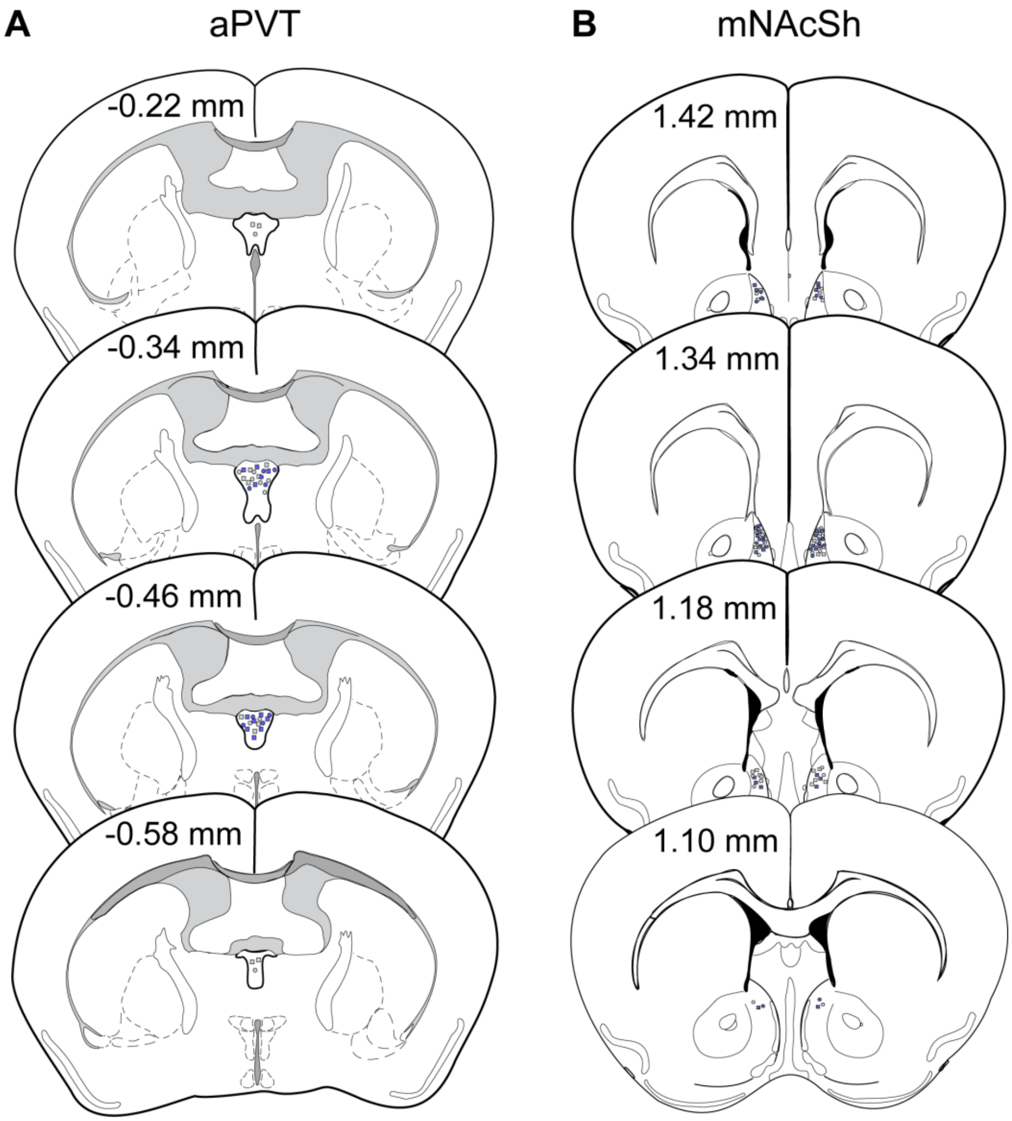


**Supplementary Fig. 2: aPVT-dorsal mNAcSh circuit eDREADD expression (related to Fig. 3). a-b)** Hit maps for DREADD virus cocktail or CON virus in aPVT (**a**) and retrograde Cre virus in mNAcSh (**b**) in DREADD (blue circles) and CON (gray circles) mice.

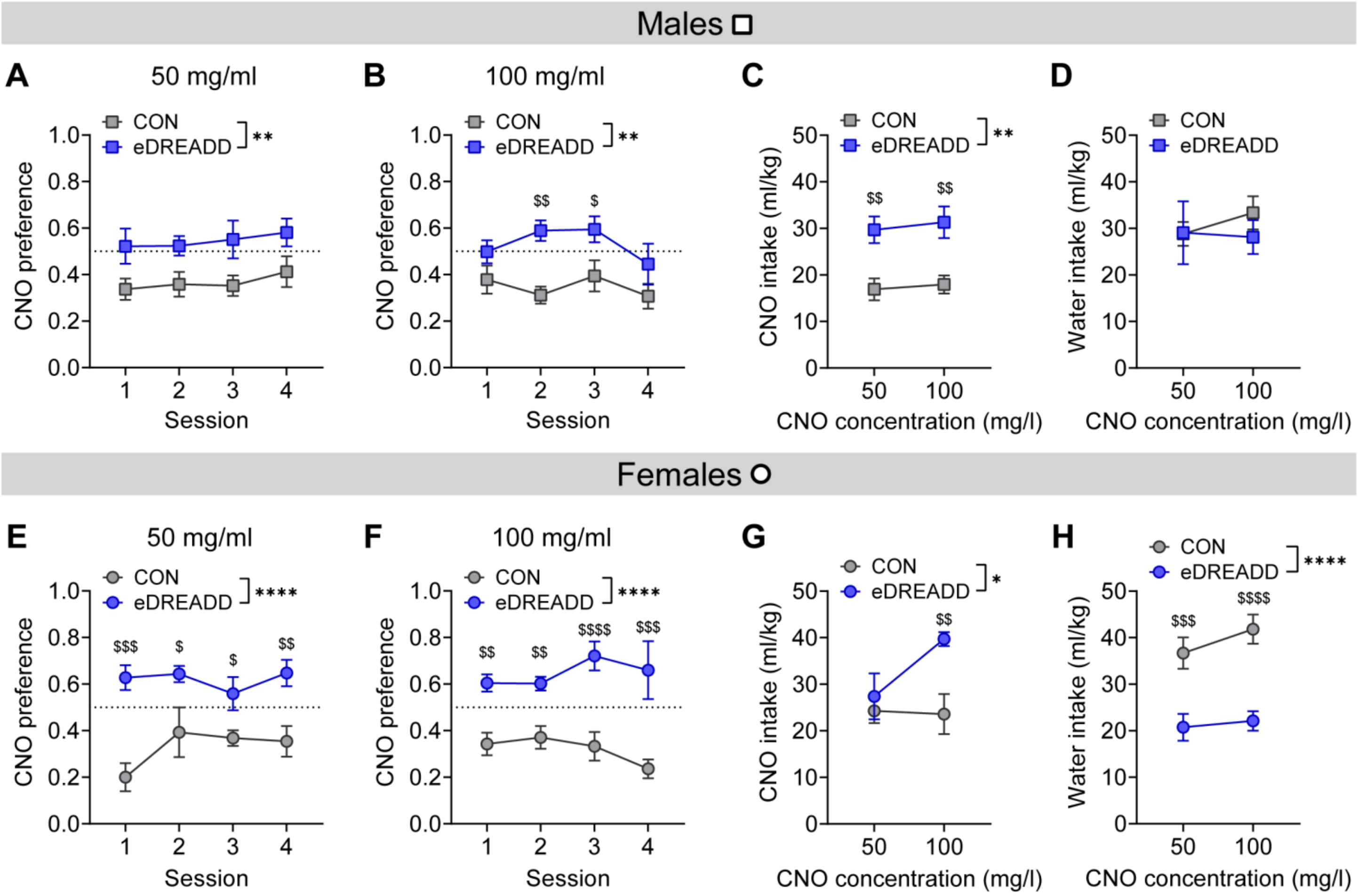


**Supplementary Fig. 3: Additional measures of CNO preference assay in aPVT-mNAcSh eDREADD mice (related to Fig. 3c-e). a-b)** CNO preference score across daily sessions with access to CNO (50 mg/l, **a**, or 100 mg/l, **b**) and water in males. **c-d)** Intake of CNO solution (**c**) and water (**d**) during the CNO preference test in males. **e-f)** CNO preference score across daily sessions with access to CNO (50 mg/l, **e**, or 100 mg/l, **f**) and water in females. **g-h)** Intake of CNO solution (**g**) and water (**h**) during the CNO preference test in females. $^{*}P < 0.05$, $^{**}P < 0.01$, $^{****}P < 0.0001$ for ANOVA main effects of DREADD as indicated; $^{\$}P < 0.05$, $^{\$\$}P < 0.01$, $^{\$\$\$}P < 0.001$, $^{\$\$\$\$}P < 0.0001$ in post hoc unpaired t-tests between CON and eDREADD mice.

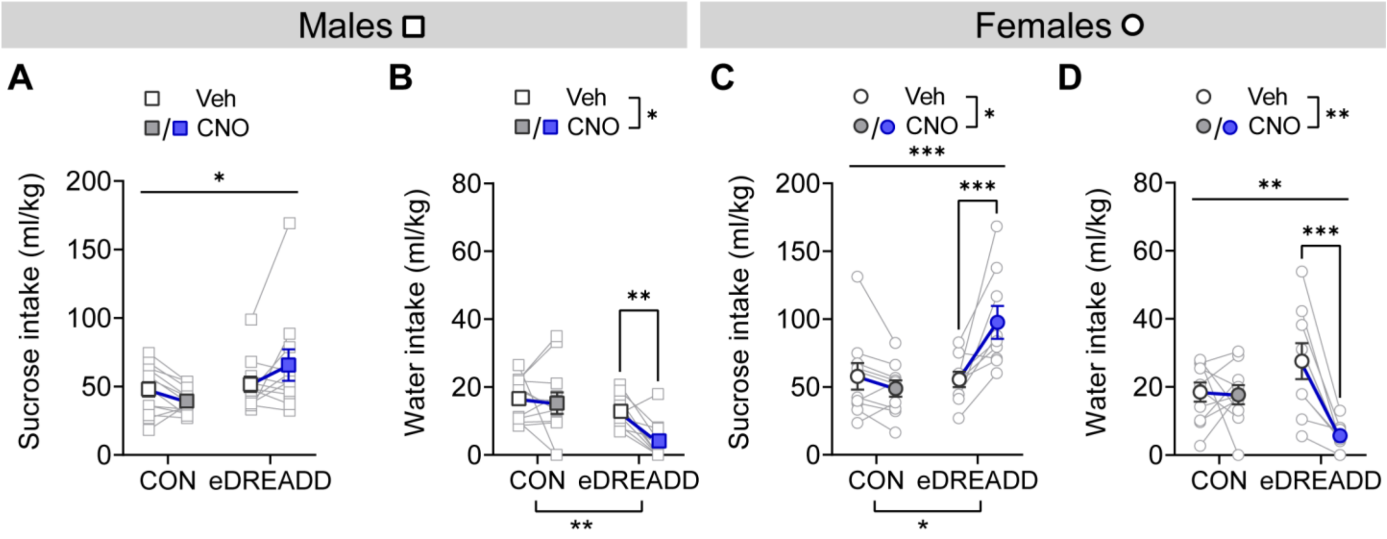


**Supplementary Fig. 4: Sucrose and water consumption during 0.5% sucrose preference test (related to Fig. 3f-h). a-b)** Intake of 0.5% sucrose (**a**) and water (**b**) during the sucrose preference test following vehicle and CNO administration in males. **c-d)** Intake of 0.5% sucrose (**c**) and water (**d**) during the sucrose preference test following vehicle and CNO administration in females. *$P < 0.05$, **$P < 0.01$, ***$P < 0.001$ for ANOVA main effects and interactions, as well as post hoc paired t-tests between Veh and CNO within group, as indicated.

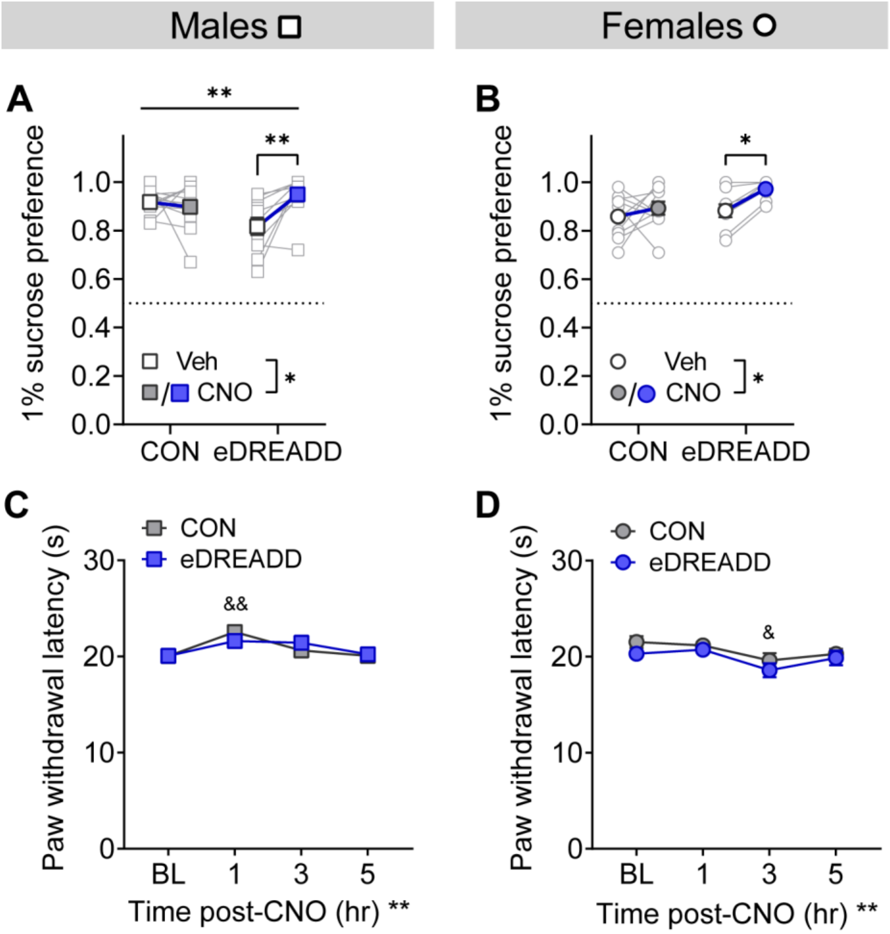


**Supplementary Fig. 5: Additional behavioral measures for aPVT-mNAcSh circuit activation (related to Fig. 3). a-b)** 1% sucrose preference was higher following CNO administration compared to vehicle (Veh) administration in eDREADD but not CON males (**a**) and females (**b**). $*P < 0.05$, $**P < 0.01$ for ANOVA main effects, interactions, and post hoc paired t-tests between Veh and CNO as indicated. **c-d)** Hot plate paw withdrawal latency at baseline (BL) and following CNO administration in males (**c**) and females (**d**). $**P < 0.01$ for ANOVA main effect of time; $^{\&}P < 0.05$, $^{\&\&}P < 0.01$ in post hoc paired t-tests between CNO time point compared to baseline (BL) in CON mice.

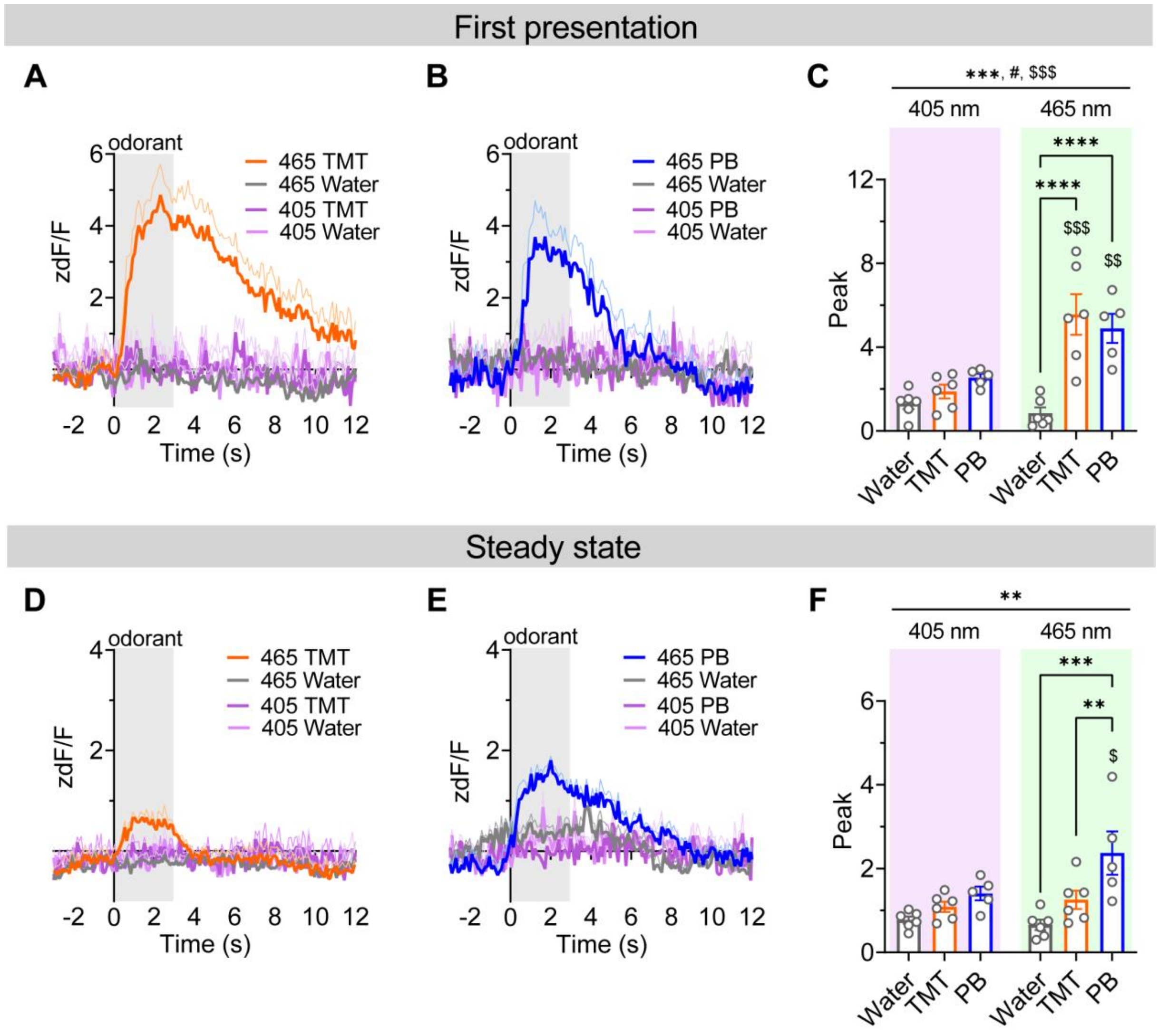


**Supplementary Fig. 6: Peak aPVT-dorsal mNAcSh terminal calcium activity during odorant presentations with control channels and conditions (related to Fig. 3). a-b)** Fiber photometry signals in the 465 nm excitation (GCaMP) and isobestic 405 nm (control) wavelength channels in aPVT synaptic terminals in the dorsal mNAcSh time-locked to the first odorant stimulus presentation on the TMT day (**a**) and PB day (**b**). **c)** Peak odorant responses from traces shown in **a,b**, showing that peak GCaMP (465 nm channel) responses to PB and TMT are higher than to water presentation and to the control 405 nm channel. **d-e)** Steady state fiber photometry signals in the 465 nm excitation (GCaMP) and isobestic 405 nm (control) wavelength channels in aPVT synaptic terminals in the dorsal mNAcSh time-locked to odorant stimulus presentations on the TMT day (**d**) and PB day (**e**). **f)** Peak odorant responses from traces shown in **d,e,** showing that peak steady state GCaMP (465 nm) responses are higher for PB than water and TMT and higher than the control 405 nm channel. Data are presented as mean +/- SEM. 465 nm traces are those from **Fig. 3d.** $^{**}P < 0.01$, $^{***}P < 0.001$, $^{****}P < 0.0001$ in 2xRM mixed model main effect of odorant and post hoc paired t-tests between odorants within channel as indicated; $^{\#}P < 0.05$ for odorant x channel interaction; $^{\$}P < 0.05$, $^{\$\$}P < 0.01$, $^{\$\$\$}P < 0.001$ in main effect of channel and post hoc paired t-tests between channel within odorant.

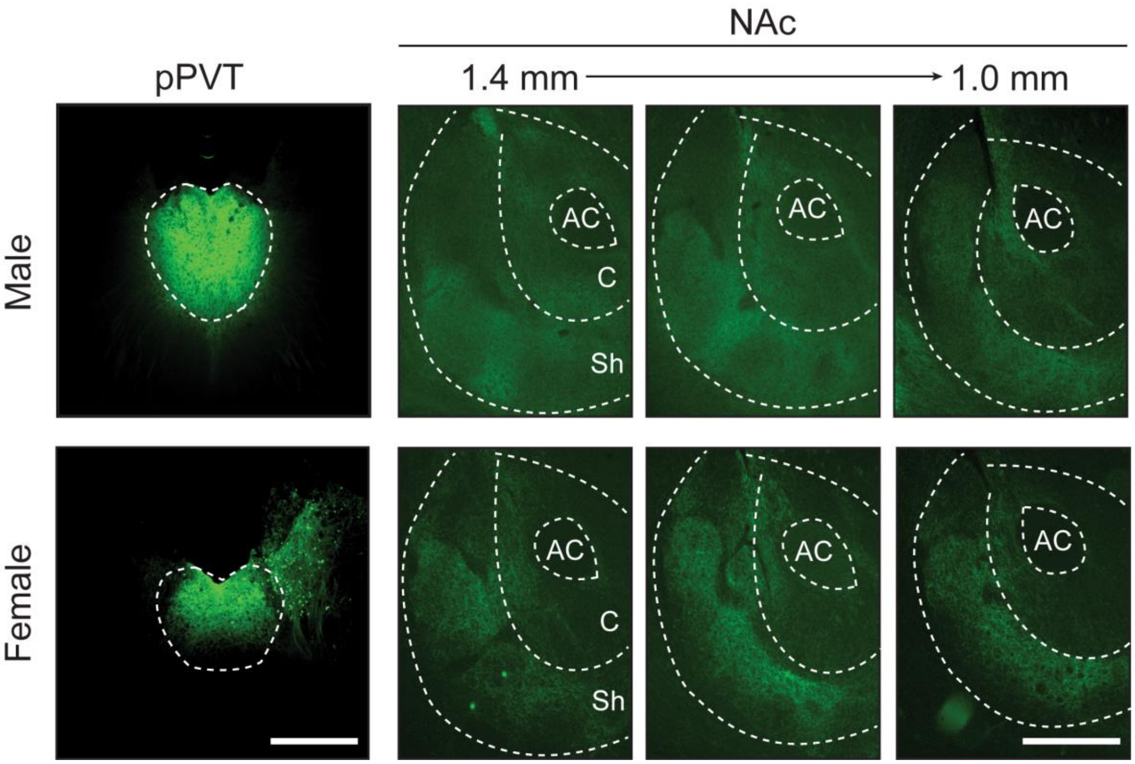


**Supplementary Fig. 7 (related to Fig. 5a): Representative images of ChR2 expression in the pPVT and across the A-P extent of the nucleus accumbens in a male (top) and female (bottom) mouse injected with ChR2 in the pPVT.** AC = anterior commissure, C and Sh = core and shell of the nucleus accumbens, respectively. Scale bars = 500 µm.

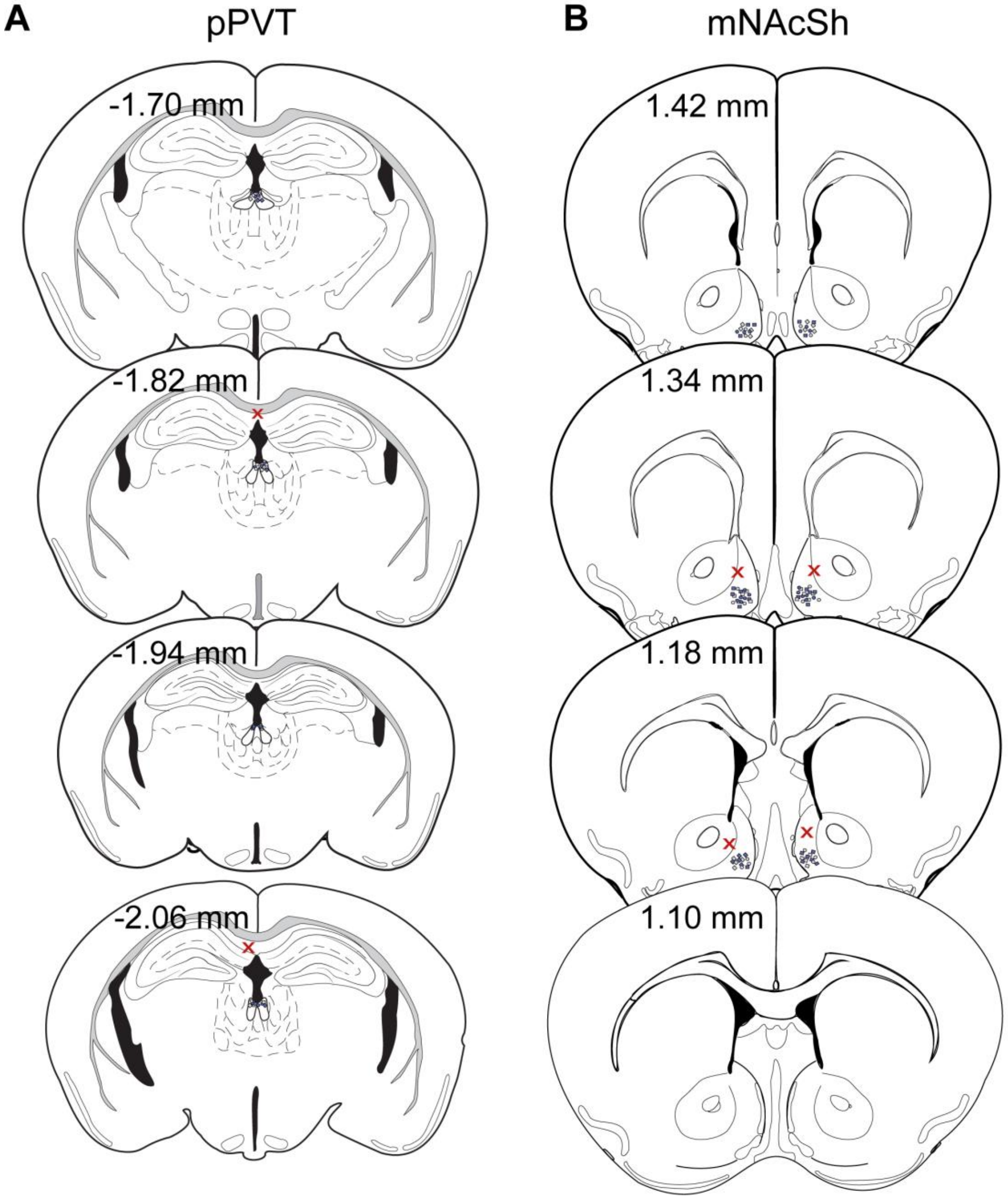


**Supplementary Fig. 8: pPVT-ventral mNAcSh circuit eDREADD expression (related to Fig. 6). a-b)** Hit maps for eDREADD/CON virus in pPVT (**a**) and retrograde Cre virus in mNAcSh (**b**), with CONs in gray and eDREADDs in blue. Red x's denote misses.

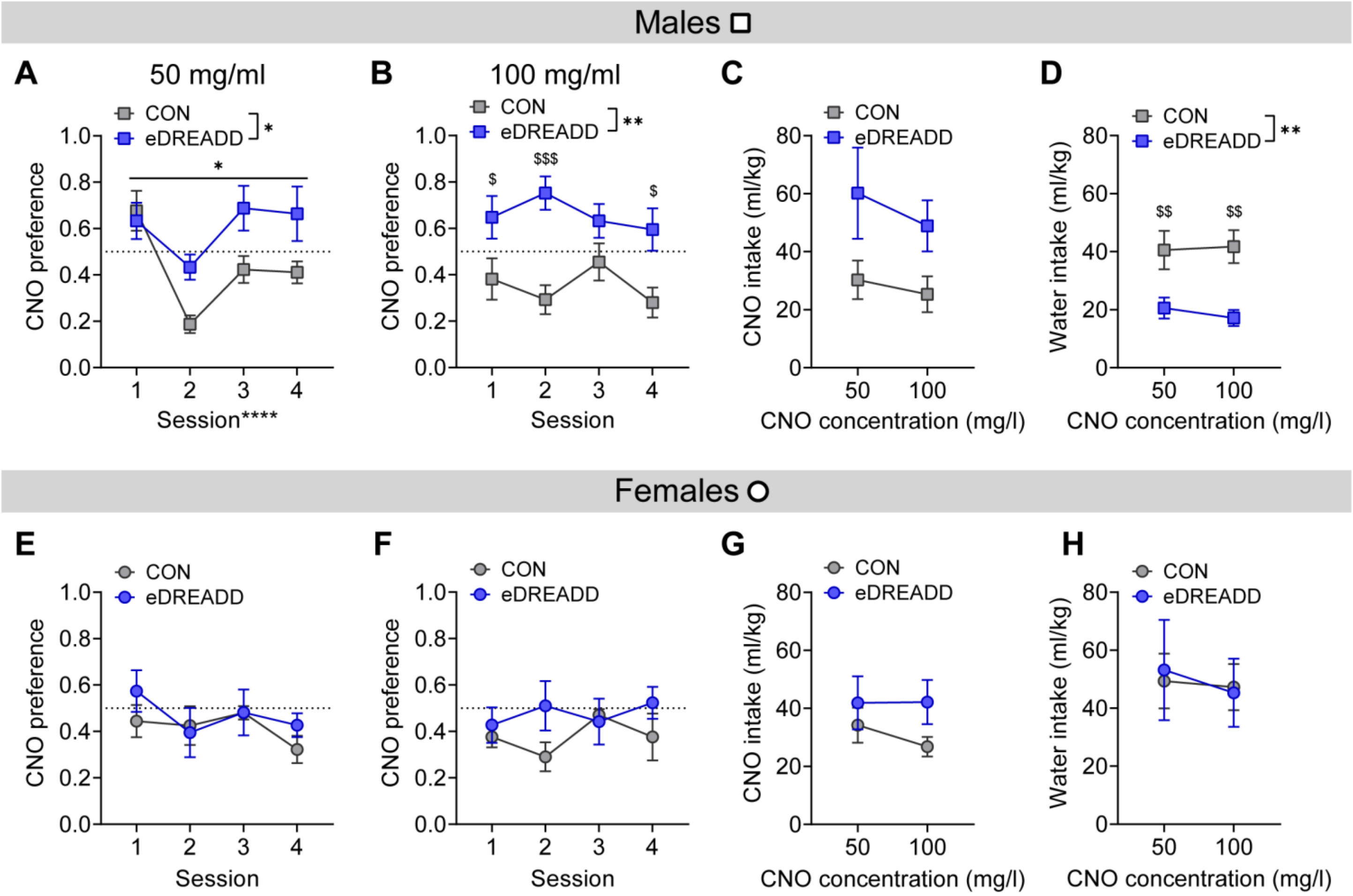


**Supplementary Fig. 9: Additional measures of CNO preference assay in pPVT-mNAcSh eDREADD mice (related to Fig. 6c-d). a-b)** CNO preference score across daily sessions with access to CNO (50 mg/l, **a**, and 100 mg/l, **b**) and water in males. **c-d)** Intake of CNO solution (**c**) and water (**d**) during the CNO preference test in males. **e-f)** CNO preference score across daily sessions with access to CNO (50 mg/l, **e**, and 100 mg/l, **f**) and water in females. **g-h)** Intake of CNO solution (**g**) and water (**h**) during the CNO preference test in females. $^{*}P < 0.05$, $^{**}P < 0.01$ for ANOVA main effects and interactions as indicated; $^{\$}P < 0.05$, $^{\$\$}P < 0.01$, $^{\$\$\$}P < 0.001$ in post hoc unpaired t-tests between CON and eDREADD mice.

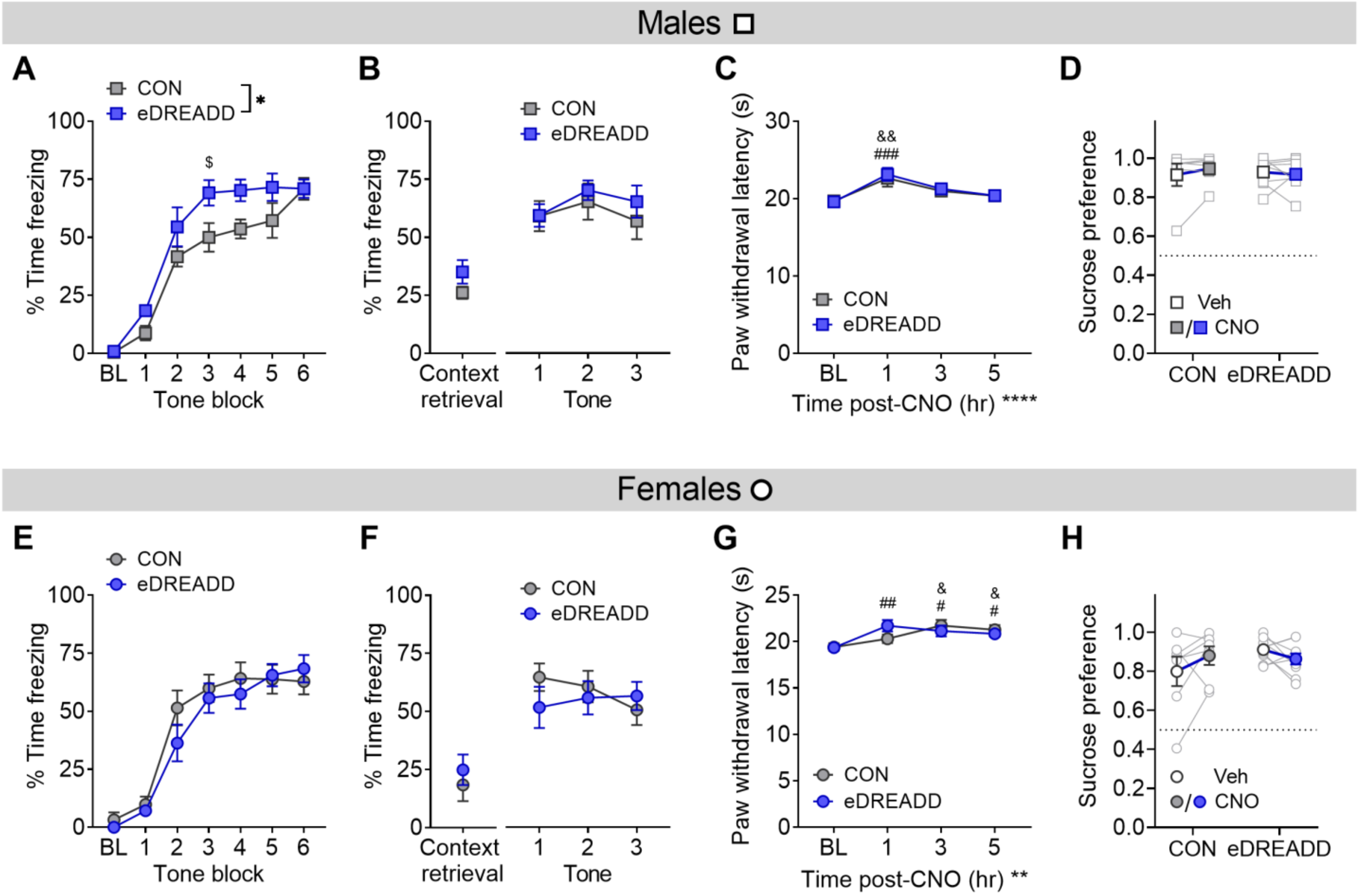


**Supplementary Fig. 10: Additional behavioral measures for pPVT-mNAcSh circuit activation (related to Fig. 6a-d). a-b)** Fear responses during Pavlovian foot shock fear conditioning (**a**) and context and cued retrieval (**b**) in males. **a)** % time freezing to tones (blocks of two tones) during acquisition on day 1 following CNO administration. **b)** % time freezing to context and tones (blocks of two tones) during retrieval on day 2 in the absence of CNO. **c)** Thermal paw withdrawal latency at baseline (BL) and following CNO administration in males. **d)** 1% sucrose preference following vehicle (Veh) and CNO administration in males. **e-f)** Fear responses during Pavlovian foot shock fear conditioning (**e**) and context and cued retrieval (**f**) in females. **e)** % time freezing to tones (blocks of two tones) during acquisition on day 1 following CNO administration. **f)** % time freezing to context and tones (blocks of two tones) during retrieval on day 2 in the absence of CNO. **g)** Thermal paw withdrawal latency at baseline (BL) and following CNO administration in females. **h)** 1% sucrose preference following vehicle (Veh) and CNO administration in females. *$P < 0.05$, **$P < 0.01$, ****$P < 0.0001$ for ANOVA main effects; $^{\$}P < 0.05$ in post hoc unpaired t-tests between CON and eDREADD mice; $^{\&}P < 0.05$, $^{\&\&}P < 0.01$ in post hoc paired t-tests between CNO time point compared to baseline (BL) in CON mice; $^{\#}P < 0.05$, $^{\#\#}P < 0.01$, $^{\#\#\#}P < 0.001$ in post hoc paired t-tests between CNO time point compared to baseline (BL) in DREADD mice.

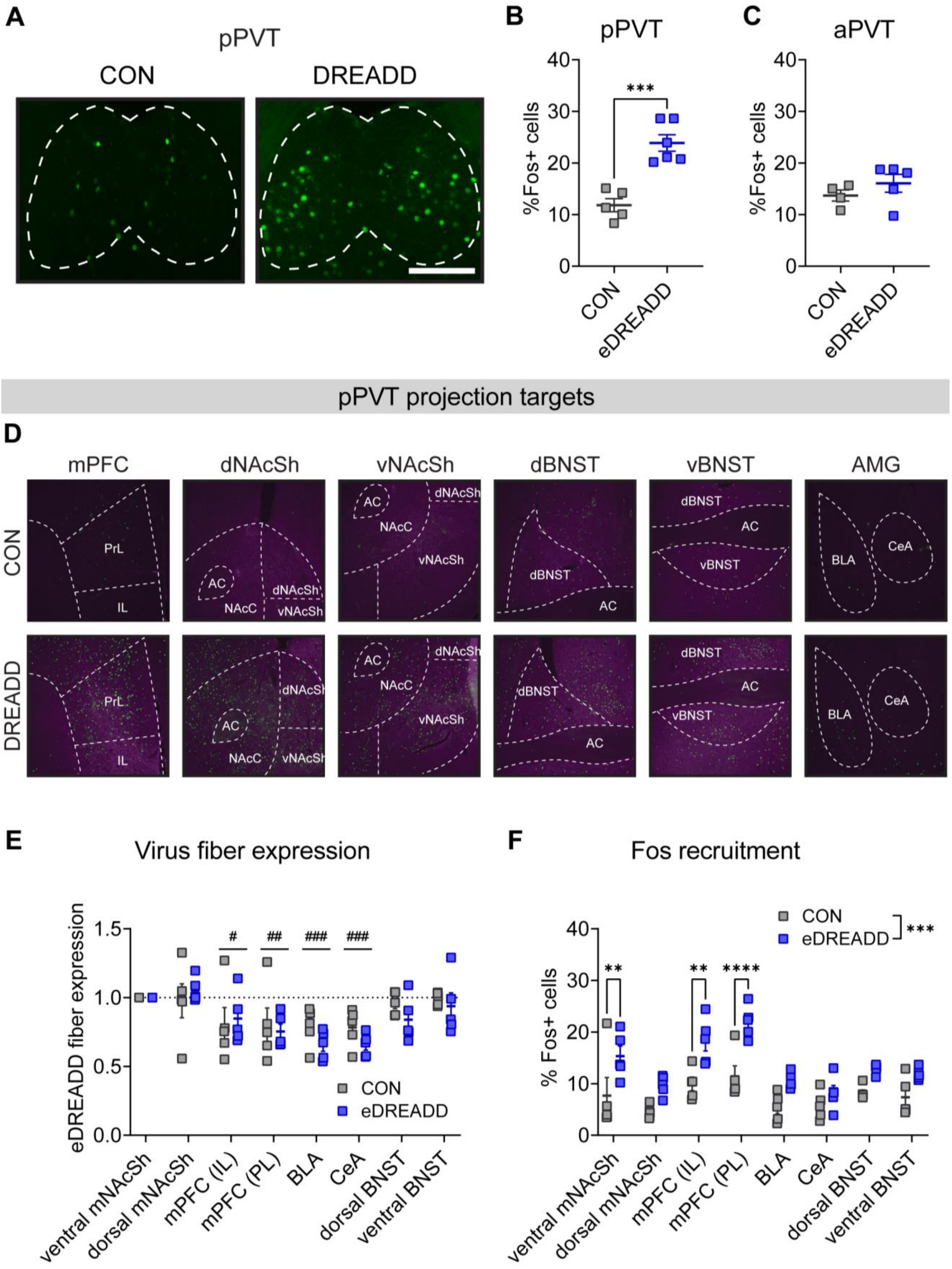


**Supplementary Fig. 11: Circuit and activity mapping following eDREADD expression and activation of the pPVT-ventral mNAcSh pathway (related to Fig. 6). a)** Representative images of Fos protein expression in the pPVT in CON and eDREADD male mice following CNO administration. Scale bar = 500 µm. **b-c)** The proportion of DAPI+ cells expressing Fos (# Fos+ cells / total DAPI) is higher in eDREADD compared to CON mice in the pPVT (**b**) but not aPVT (**c**). **d)** Representative images of eDREADD and Fos expression in known projection targets of the pPVT in mouse with eDREADD expression in the pPVT-ventral mNAcSh circuit. **e)** eDREADD and CON virus expression in target regions normalized to expression in the ventral mNAcSh, showing that regions outside the mNAcSh and BNST show lower terminal expression than ventral mNAcSh. **f)** Following CNO activation of the pPVT-ventral mNAcSh eDREADD, Fos expression was higher in the ventral mNAcSh and infralimbic (IL) and prelimbic (PL) subregions of the medial PFC compared to CON mice. $^{**}P < 0.01$, $^{***}P < 0.001$, $^{****}P < 0.0001$ for ANOVA main effects and post hoc and a priori unpaired t-tests as indicated. $^{\#}P < 0.05$, $^{\#\#}P < 0.01$, $^{\#\#\#}P < 0.001$ in post hoc paired t-tests between virus expression compared to ventral mNAcSh for regions indicated.

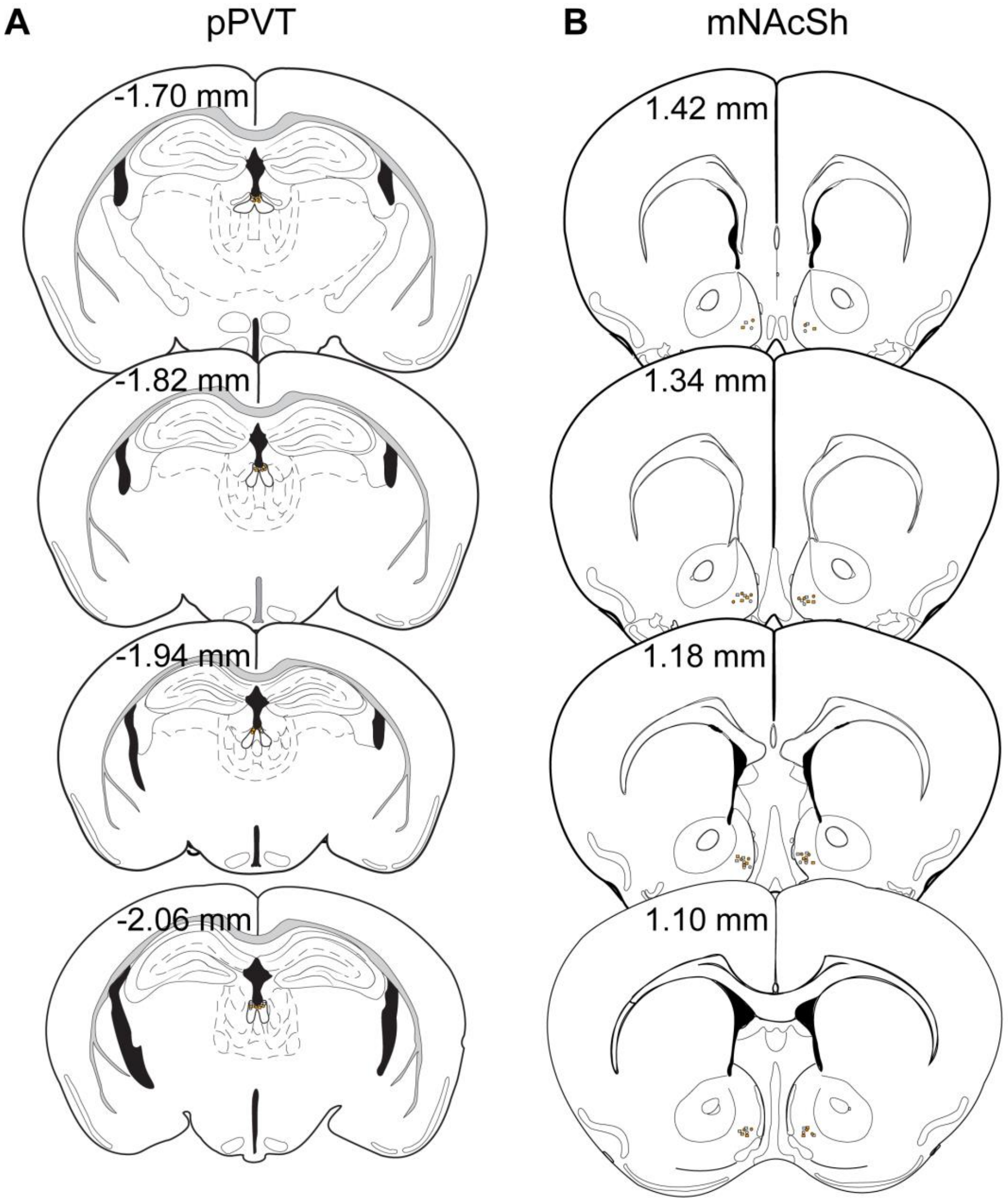


**Supplementary Fig. 12: pPVT-ventral mNAcSh circuit iDREADD expression (related to Fig. 7). a-b)** Hit maps for iDREADD/CON virus in pPVT (**a**) and retrograde Cre virus in ventral mNAcSh (**b**) in CON (gray circles) and iDREADD (orange circles) mice.

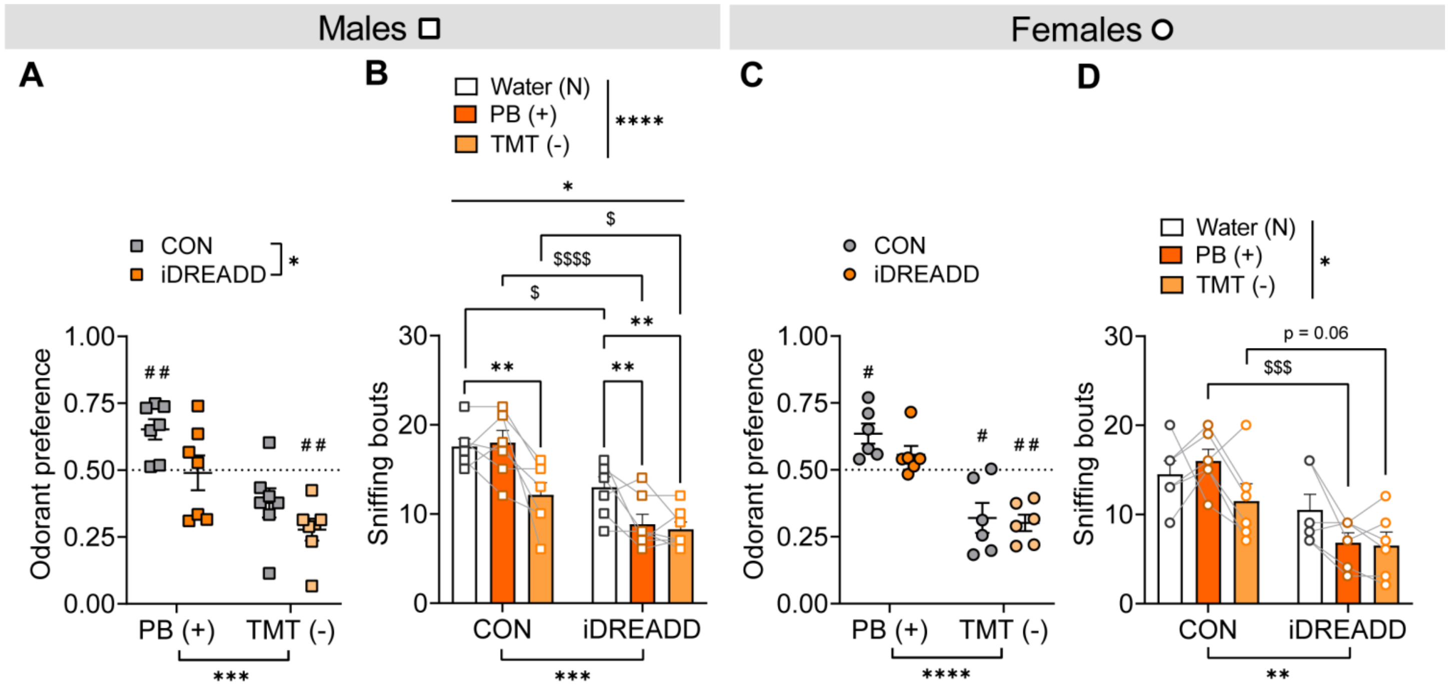


**Supplementary Fig. 13: Additional measures for chemogenetic inhibition of the pPVT- ventral mNAcSh circuit during behavior (related to Fig. 7). a-b)** Odorant preference scores for positively and negatively valenced odorants over water (**a**) and number of bouts of odorant sniffing (**b**) following CNO administration in males. iDREADD mice displayed reduced investigation of all odorants compared to CONs. **c-d)** Odorant preference scores (**c**) and number of bouts of odorant sniffing (**d**) following CNO administration in females. $^{*}P < 0.05$, $^{**}P < 0.01$, $^{***}P < 0.001$, $^{****}P < 0.0001$ for ANOVA main effects, interactions, and post hoc t-tests as indicated; $^{\$}P < 0.05$, $^{\$\$\$}P < 0.001$, $^{\$\$\$\$}P < 0.0001$ in post hoc t-test between CON and iDREADD mice. $^{\#}P < 0.05$, $^{\#\#}P < 0.01$ for one-sample t-tests compared to 0.50 for odorant preference within each group.
